# A clusterability measure for single-cell transcriptomics reveals phenotypic subpopulations

**DOI:** 10.1101/2021.05.11.443685

**Authors:** Maria Mircea, Mazène Hochane, Xueying Fan, Susana M. Chuva de Sousa Lopes, Diego Garlaschelli, Stefan Semrau

**Affiliations:** Leiden Institute of Physics, Leiden University, Leiden, The Netherlands; Leiden Academic Center for Drug Research, Leiden University, Leiden, The Netherlands; Department of Anatomy and Embryology, Leiden University Medical Center, Leiden, The Netherlands; Networks Unit, IMT School for Advanced Studies, Lucca, Italy

## Abstract

The ability to discover new cell populations by unsupervised clustering of single-cell transcriptomics data has revolutionized biology. Currently, there is no principled way to decide, whether a cluster of cells contains meaningful subpopulations that should be further resolved. Here we present SIGMA, a clusterability measure derived from random matrix theory, that can be used to identify cell clusters with non-random sub-structure, testably leading to the discovery of previously overlooked phenotypes.

## Main

Unsupervised clustering methods^1–4^ are integral to most single-cell RNA-sequencing (scRNA-seq) analysis pipelines^5^. All existing clustering algorithms have adjustable parameters, which have to be chosen carefully to reveal the true biological structure of the data. If the data is over-clustered, many clusters are driven purely by technical noise and do not reflect distinct biological states. If the data is under-clustered, subtly distinct phenotypes might be grouped with others and will thus be overlooked. Existing tools to assess clustering quality, such as the widely used silhouette coefficient, cannot reveal if the variability within a cluster is due to the presence of subpopulations or random noise.

To alleviate this problem, we developed SIGnal-Measurement-Angle (SIGMA), a clusterability measure for scRNA-seq data. We consider clusterability to be the theoretically achievable agreement with the unknown ground truth clustering, for a given signal-to-noise ratio. Importantly, our measure can estimate the level of achievable agreement without knowledge of the ground truth. High clusterability (indicated by SIGMA close to 1) means that multiple phenotypic subpopulations are present and clustering algorithms should be able to distinguish them. Low clusterability (indicated by SIGMA close to 0) means that the noise is too strong for even the best possible clustering algorithm to find any clusters accurately. If SIGMA equals 0, the observed variability within a cluster is consistent with random noise.

To derive SIGMA, we considered the unobserved, actual gene expression profiles (the signal matrix) as a perturbation to a random noise matrix (Fig. 1a). This is the exact opposite of the conventional view, which considers noise as a perturbation to a signal. Note that both the biological variability within a phenotype as well as technical variability (due to variable capture and conversion efficiencies etc.) contribute to random noise. Our point of view allowed us to leverage well-established results from random matrix theory^6,7^ and perturbation theory^8^. Briefly, we first calculate the singular value distribution of the measured expression matrix. If the data is preprocessed appropriately (Extended Data Fig. 1), the bulk of this distribution is described by the Marchenko-Pastur (MP) distribution, which corresponds to the random component of the measurement. The singular values outside of the MP distribution and above the Tracy Widom (TW) threshold correspond to the signal (i.e. the unobserved gene expression profiles). Using just these singular values and the dimensions of the measurement matrix, we can calculate the angles between the singular vectors of the measured expression matrix and those of the (unobserved) signal matrix. SIGMA is the squared cosine of the smallest angle. See Supplementary Note 1 for a detailed derivation. Simulations of data sets with varying signal-to-noise ratios illustrate the calculation of SIGMA (Fig. 1b,c). Data sets with higher signal-to-noise ratios have more easily separable clusters and larger singular values outside of the MP distribution (Fig. 1b). By definition, that results in higher values of SIGMA (Fig. 1c).

**Fig. 1.**
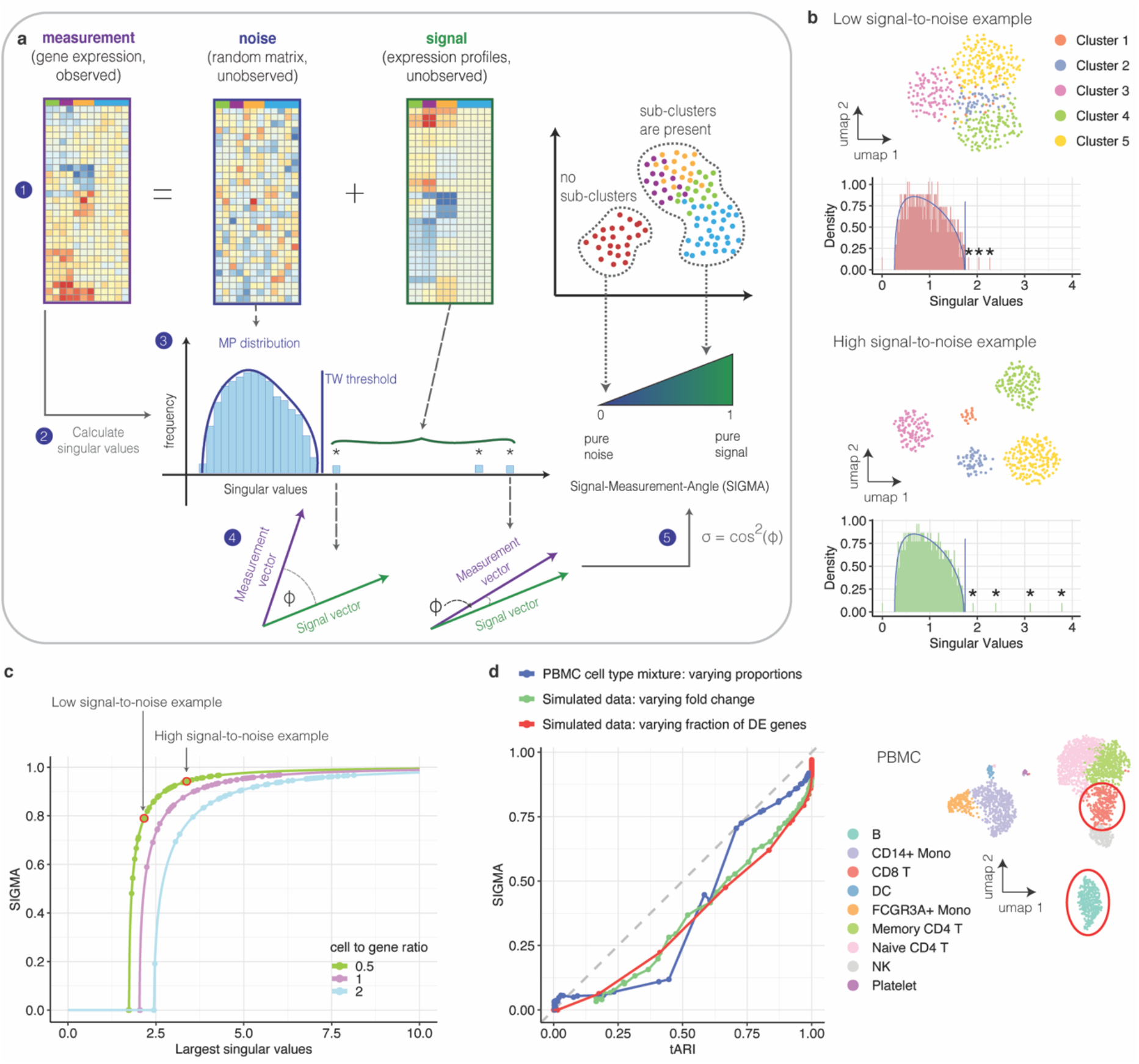
SIGMA, a clusterability measure for scRNA-seq data derived from random matrix theory, is a proxy for the theoretically achievable adjusted rand index (tARI). **a** Scheme illustrating the rationale. **b** Singular value spectra of simulated data sets with 5 clusters and different levels of noise; Red: low signal-to-noise, Green: high signal-to-noise. The MP distribution is indicated by a solid line, significant singular values are highlighted with asterisks. Inserts show UMAPs of the data. The data set with a higher signal-to-noise ratio has more significant singular values and those singular values are bigger. **c** Value of the largest singular value versus SIGMA for simulated data. Arrows indicate where the examples from panel a are located. The relationship between the largest singular values and SIGMA only depends on the dimensions of the expression matrix. Simulations with different cell-to-gene ratios are shown in different colors. **d** SIGMA versus theoretically achievable ARI (tARI). Red data points: Simulated data sets with two clusters. The number of differentially expressed (DE) genes was varied, the log fold change between clusters was fixed. Green data points: Simulated data sets with two clusters. The log fold change between clusters was varied, the number of differentially expressed genes was fixed. Blue data points: Two synthetic clusters were created by weighted averages of cells from two clusters in the PBMC data set. Cluster weights were varied. The grey dashed line indicates identity. Inset: UMAP of PBMC data set with the two clusters used indicated by red solid circles.

To show that SIGMA is a proxy for clusterability, we have to make the concept of clusterability more precise and quantifiable. First, we adopted the Adjusted Rand Index (ARI)^9^ as a well-established measure for the agreement between an empirically obtained clustering and the ground truth. Next, we argued that perfect agreement with the ground truth (ARI = 1) is not achievable in the presence of noise, even with the best conceivable clustering algorithm. Using a simple case of two clusters of cells with varying signal-to-noise ratios, we estimated the Bayesian error rate^10^ (i.e. the lowest possible error) for this binary classification problem in simulated data (Extended Data Fig. 2a). Based on this error rate, we calculated a theoretically achievable ARI (tARI, see also Supplementary Note 1). We showed empirically that commonly used clustering methods do not exceed this limit (Extended Data Fig. 2b,c). The tARI, therefore, quantifies our notion of clusterability. Importantly, SIGMA is strongly correlated with the tARI (Fig. 1d) and thus allows us to estimate clusterability without knowing the ground truth. To confirm this result in experimentally measured data, we chose two very distinct clusters from a PBMC data set^11^ and created two new clusters as weighted averages, which allowed us control over the signal-to-noise ratio. Also for this data, SIGMA strongly correlates with the tARI (Fig. 1d). As an alternative to the tARI, we also calculated the theoretically achievable silhouette coefficient^12^ (tSIL), which considers the distances between the best possible clusters (Extended Data Fig. 3 a-c). The tSIL quickly jumps to higher values for minimal deviations from pure noise, due to the correct classification of a few outlier cells, which makes it less useful for assessing overall clusterability. We also compared SIGMA to ROGUE^13^, a recently published clusterability measure (Extended Data Fig. 3d). In contrast to SIGMA, ROGUE does not show collinearity with the tARI. Therefore, ROGUE seems to estimate a notion of clusterability that is distinct from our point of view.

To further characterize the performance of SIGMA on experimental data sets with known ground truth, we used a measurement of purified RNA from 3 cell types, mixed at different ratios^14^ (Extended Data Fig. 4a). We noticed that the amount of input RNA used for each mixture was a confounding factor that influenced the value of SIGMA (Extended Data Fig. 4b,c). It is well-established that various factors drive artefactual variability in single-cell RNA-seq data^11,15^. We therefore introduced a regression step, that removes the influence of any nuisance variables, such as the number of total counts per cell, ribosomal gene expression, mitochondrial gene expression or cell cycle phase (Extended Data Fig. 4b-c, see also Supplementary Note 1). After correction, SIGMA successfully indicated the presence or absence of sub-clusters for all tested combinations of the 7 original RNA mixtures (Extended Data Fig. 5). By contrast, ROGUE only indicated the presence of sub-structure when the merged clusters were very clearly distinguishable (Extended Data Fig. 5b,c). This indicates that SIGMA is a more sensitive measure, which detects differences between highly similar phenotypes.

In full analogy to the reasoning outlined so far, our approach can also be used to characterize variability in the space of genes. We call this conjugate measure G-SIGMA (see Supplementary Note 1 for the derivation). Data sets with higher signal-to-noise ratios are characterized by higher values of G-SIGMA (Extended Data Fig. 6a), which indicates a more accurate estimation of differential gene expression after sub-clustering. Furthermore, genes with higher absolute values in a certain gene-singular vector drive the variability observed in the corresponding cell-singular vector (Extended Data Fig. 6 b-d). Our approach thus not only identifies relevant sub-structure in a cell cluster but can also reveal the genes responsible for it. This is not a direct replacement for differential expression tests, but a way to understand the variability within the cell-singular vectors.

Finally, we tested the performance of SIGMA and G-SIGMA in measurements of complex tissues. In a data set of bone marrow mononuclear cells (BMNC)^16^ we calculated SIGMA for the clusters reported by the authors. After correction for confounding factors (Extended Data Fig. 4 d,e), SIGMA corresponded well with a visual inspection of the cluster UMAPs (Fig. 2a). For all clusters, the bulk of the singular value distribution was well-described by the MP distribution and, by construction, only clusters with SIGMA > 0 had significant singular values (Fig. 2b). Reassuringly, many progenitor cell types received a high SIGMA (indicating possible sub-structure) in agreement with the known higher variability in these cell types. Ranking existing clusters by G-SIGMA resulted in a very similar order (Extended Data Fig. 7a). To confirm the presence of relevant sub-structure, we sub-clustered the two original clusters with the highest SIGMA (Extended Data Fig. 7 b-e). In the red blood cell (RBC) progenitors, we identified 4 subsets that correspond to different stages of differentiation, ranging from erythroid precursors to highly differentiated RBCs as identified by F.V Mello et al.^17^. In the dendritic cell (DC) progenitor cluster, two sub-clusters were identified, which correspond to precursors of either classical or plasmacytoid DCs^18^. For both examples, the variance-driving genes found in the gene-singular vectors were localized to their corresponding clusters (Extended Data Fig. 7 c,d) and overlapped strongly with differentially expressed genes found after sub-clustering (see Supplementary Table 2).

**Fig. 2.**
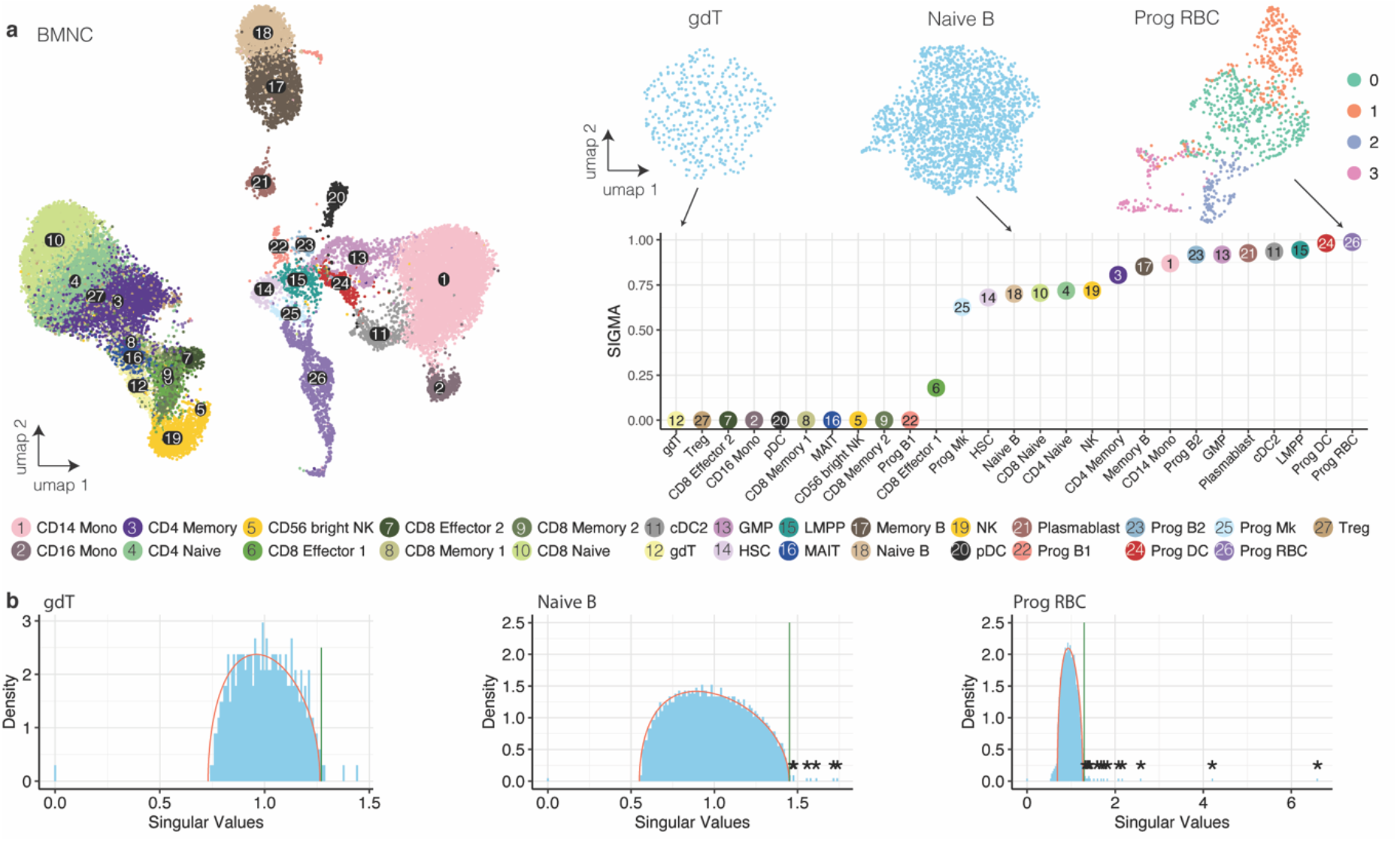
Application of SIGMA to BMNC data can drive the discovery of biologically meaningful sub-clusters. **a** UMAP and SIGMA for BMNC data set. Inset: UMAP of clusters with low, intermediate, and high values of SIGMA. **b** MP distribution of clusters with low, intermediate, and high values of SIGMA.

In a second example, we applied SIGMA to a fetal human kidney data set we published previously^19^ (Fig 3a). As for BMNCs, SIGMA corresponded well with a qualitative assessment of cluster variability and G-SIGMA resulted in a similar ranking (Extended Data Fig. 8a). Sub-clustering of the cluster with the highest SIGMA, ureteric bud/collecting duct (UBCD), revealed a subset of cells with markers of urothelial cells (*UPK1A, KRT7*) (Fig. 3b, Extended Data Fig. 8b-e). Immunostaining of these two genes, together with CDH1 expressed in the collecting system, in week 15 fetal kidney sections confirmed the presence of the urothelial subcluster (Fig. 3c, Extended Data Fig. 9a). Another subset of cells we did not find in our original analysis, are the parietal epithelial cells (PECs), which could now be identified within the SSBpr cluster (S-shaped body proximal precursor cells) (Fig. 3b, Extended Data Fig. 8b-e). To reveal these cells *in situ*, we stained for AKAP12 and CAV2, which were among the top differentially expressed genes in this subcluster (Supplementary Table 3), together with CLDN1, a known marker of PECs, and MAFB, a marker of the neighboring podocytes (Fig. 3d, Extended Data Fig. 9b). Together with the PECs and proximal tubule precursor cells, SSBpr also contained a few cells that were misclassified in the original analysis, indicating the additional usefulness of SIGMA as a means to identify clustering errors. Further analysis of a cluster of interstitial cells (ICa) revealed multiple subpopulations (Fig. 3b, Extended Data Fig. 8b-e). Immunostainings revealed that a POSTN-positive population is found mostly in the cortex, often surrounding blood vessels, whereas a SULT1E1-positive population is located in the inner medulla and papilla, often surrounding tubules (Fig. 3e, Extended Data Fig. 9c). CLDN11, another gene identified by analysis of the gene-singular vectors (Extended Data Fig. 8b-e) was found mostly in the medulla, but also in the outermost cortex. A more detailed, biological interpretation of the results can be found in Supplementary Note 2.

**Fig. 3.**
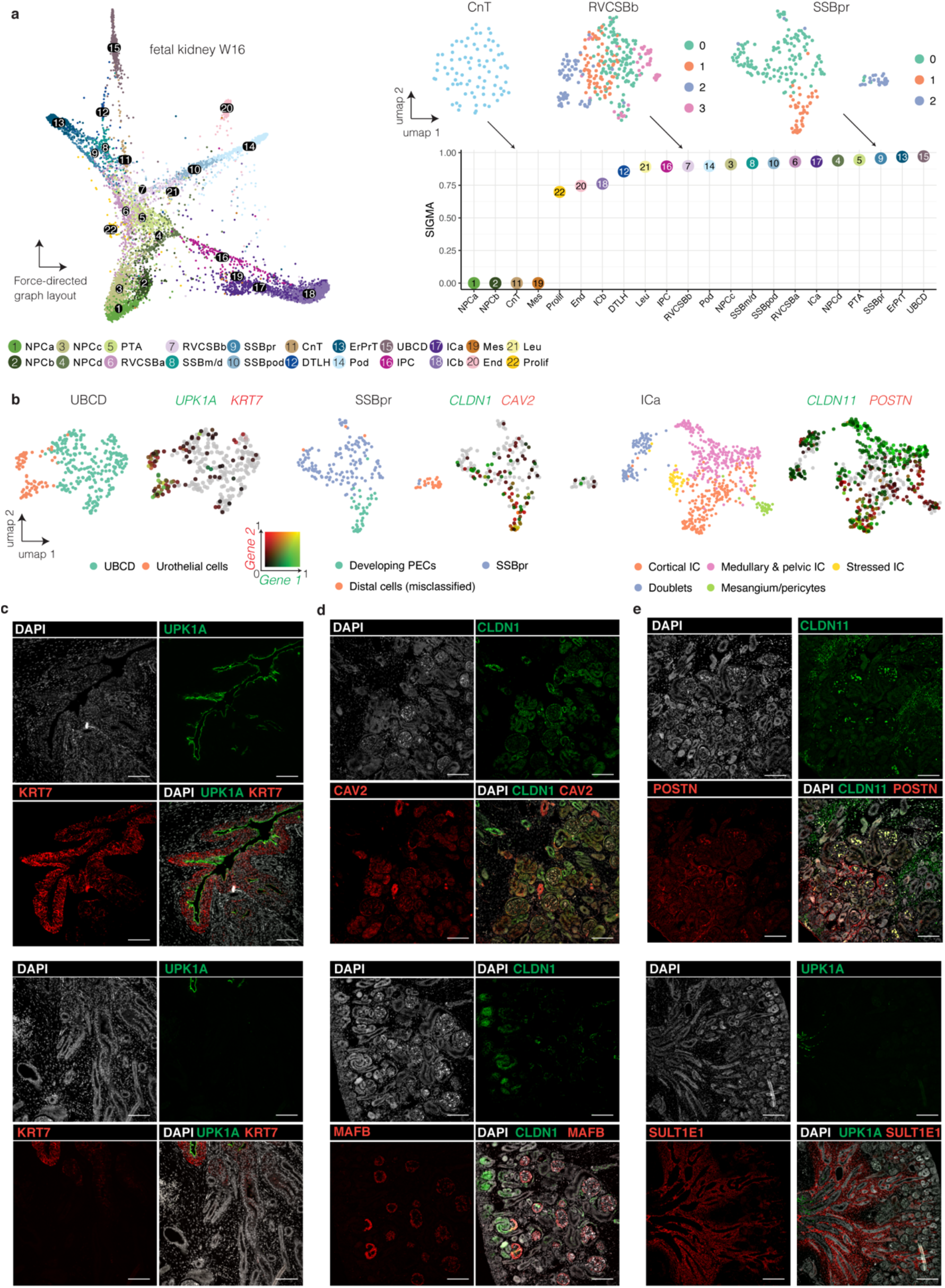
Application of SIGMA to a human fetal kidney leads to the discovery of biologically meaningful sub-clusters. **a** Force-directed graph layout and SIGMA for the fetal kidney data set. Inset: UMAP of clusters with low, intermediate, and high values for SIGMA. **b** UMAPs of the UBCD, SSBpr, and ICa clusters. Left: Colors indicate sub-clusters. Right: Colors indicate the log-normalized gene expression of the two indicated genes. One gene follows the red color spectrum, the other gene the green color spectrum. The combined expression of two genes is either dark (low expression in both genes) or yellow (high expression in both genes). **c-e** Immunostainings of week 15 fetal kidney sections. **c** UPK1A and KRT7 are expressed in the urothelial cells of the developing ureter (upper panel) and absent from the tubules in the adjacent inner medulla (lower panel). **d** PECs express CLDN1 and CAV2 (upper panel), as well as CLDN1 at the capillary loop stage and further (lower panel). MAFB staining is found in podocytes and their precursors in the SSB (lower panel). **e** CLDN11 and POSTN are expressed in interstitial cells visualized by immunostaining (upper panel), expression of SULT1E1 in the interstitial cells surrounding the ureter (UPK1A), and the tubule in the inner medulla (lower panel). Scale bars: 100 µm.

In summary, we presented SIGMA, a clusterability measure that can help to detect easily overlooked, subtle phenotypes in scRNA-seq data. Our approach also identifies variance-driving genes and brings renewed awareness to random noise as a factor setting hard limits on clustering and identifying differential expression.

## Methods

### Preprocessing

Before applying the method to simulated or measured single-cell RNA-seq data sets, several preprocessing steps are necessary. The raw counts are first normalized and log-transformed. Next, the expression matrix is standardized, first gene-wise, then cell-wise. These steps assure the proper agreement of the bulk of the singular value distribution with the MP distribution (Extended Data Fig. 1). See also Supplementary Note, Section 3.1.

### Signal-Measurement angle (SIGMA)

SIGMA is based on the assumption that the expression matrix 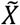 measured by scRNA-seq, can be written as the sum of a random matrix *X*, which contains random biological variability and technical noise, and a signal matrix *P*, which contains the unobserved expression profiles of each cell:

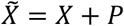

Note that in this decomposition, cells that belong to the same cell type have identical expression profiles in the signal matrix *P*. This notion of clusterability, based on the signal-to-noise ratio, is inspired by the notion of detectability in networks^20,21^.

Treating the signal matrix *P* as a perturbation to the random matrix *X*, we can apply results from both random matrix theory and low-rank perturbation theory. Random matrix theory^22,23^ predicts that the singular value distribution of *X* is a Marchenko-Pastur (MP) distribution^7,24,25^, which coincides with the bulk of the singular value distribution^6,26,27^ of 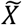. The singular values of 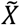 above the values predicted by the MP distribution characterize the signal matrix *P*. Since the agreement with the MP distribution holds strictly only for infinite matrices, we use two additional concepts to identify relevant singular values exceeding the range defined by the MP distribution. The Tracy-Widom^25,28^ (TW) distribution describes the probability of a singular value to exceed the MP distribution, if the matrix is finite. Additionally, since singular vectors of a random matrix are normally distributed, relevant singular vectors have to be significantly different from normal^6^. To test for normality we used the Shapiro-Wilk test.

We apply low-rank perturbation theory^8^ to calculate the singular values (*θ*_*i*_) of *P* from the relevant singular values (γ_*i*_) of the measured expression matrix 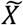:

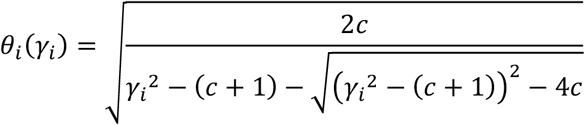

where c is the cell-to-gene ratio, i.e. the total number of cells divided by the total number of genes.

The values of *θ*_*i*_ are then used to obtain the angles ϕ_*i*_ between the singular vectors of 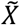 and *P*. These angles are conveniently expressed in terms of their squared cosine as

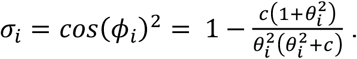

The squared cosine of the smallest angle, i.e. the largest squared cosine, is then used as a measure of clusterability:

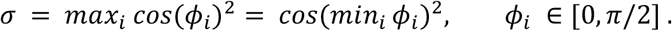

For a detailed derivation of SIGMA, see Supplementary Note 1, Section 2.1-2.4.

### Confounder regression

scRNA-seq data contains various confounding factors that drive uninformative variability. These either emerge from technical issues (such as the varying efficiency of transcript recovery, which cannot be fully eliminated by normalization) or biological factors (such as cell cycle phase, metabolic state, or stress), see Extended Data Fig. 4. To account for these factors, a regression step, inspired by current gene expression normalization methods^11,15^, is included. If a singular vector is biased by any of the considered confounders, its singular value will be reduced, which leads to a lower SIGMA value. See also Supplementary Note, Section 3.2.

### Theoretically achievable clustering quality

To construct a Bayes classifier^10^, which achieves the minimal error rate, we need to know the ground truth clustering. Hence, we used data simulated with Splatter^29^, containing two clusters. For each ground truth cluster, we fit a multidimensional Gaussian to the corresponding elements of the singular vectors (see Extended Data Fig. 2a). We only consider singular vectors with singular values larger than predicted by the MP distribution. For the fit, we use the mclust^30^ R package (V 5.4.6). We then construct a classifier by assigning a cell to the cluster for which it has the highest value of the fitted Gaussian distribution. This classifier is thus approximately a Bayes classifier (for a true Bayes classifier, we would need to know the exact distributions of the singular vector entries). The ARI^9^ calculated based on this classification is thus approximately the best theoretically achievable ARI (tARI).

The silhouette coefficient^12^ was calculated on Euclidean distances in the first singular vectors and the average silhouette coefficient was reported. In the RNA-mix data, Euclidean distances were calculated using singular vectors whose singular values exceed the range defined by the MP distribution and the ground truth clustering. For the simulated data sets with 2 clusters, the silhouette coefficient was calculated on the first singular vector and the clusterings produced by the different methods (Extended Data Fig. 3). tSIL was calculated with Euclidean distances in the first singular vector on the best theoretically achievable clustering. The calculation of tARI and tSIL is described in more detail in Supplementary Note 1, section 2.5.

### Clustering methods

For the validation of the tARI and tSIL, several clustering methods were used on simulated data with two clusters. Seurat clustering^1^ was performed with the *Seurat R package* with 10 principal components (PCs) and 20 nearest neighbors. Three different resolution parameters were used: 0.1, 0.6, and 1.6. Scanpy clustering^2^ was performed with the *scanpy python package* with 10 PCs and 20 nearest neighbors. Three different resolution parameters were used: 0.1, 0.6, and 1.6. Hierarchical clustering^4^ was performed on the first 10 PCs and Euclidean distances. The hierarchical tree was built with the Ward linkage and the tree was cut at a height where 2 clusters could be identified. K-means^3^ was performed on the first 10 PCs using Euclidean distances and two centers. TSCAN^31^ was calculated on the first 10 PCs.

### ROGUE

ROGUE^13^ is an entropy-based clusterability measure. A null model is defined under the assumption of Gamma-Poisson distributed gene expression and its differential entropy is then compared to the actual differential entropy of the gene expression.

For the RNA-mix data set ROGUE (V 1.0) was used with 1 sample (see Fig S5), “UMI” platform, and a span of 0.6. For the simulated data sets, ROGUE was used with k = 10 (Extended Data Fig. 2 d).

### Variance driving genes

Genes that drive the variance in the significant singular vectors can be used to explore the biological information in the sub-structures. Genes with large positive or negative entries in a gene-singular vector are localized in cells with high positive or negative entries in the corresponding cell-singular vector. It is also possible to assess the signal-to-noise ratio for the genes by calculating the angle between the gene singular vectors of the measured expression matrix 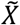 and the gene singular vectors of the signal matrix *P*, given by^15^

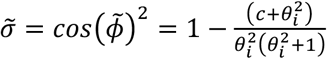

where c is the cell-to-gene ratio. We call 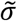 the gene SIGMA (G-SIGMA). See Supplementary Note 1, section 2.4 for a more detailed discussion.

### Data sets

Simulated data were produced with the *splatter*^29^ *R package (V 1*.*10*.*1)*. The parameters used for the simulation are shown in Supplementary Table 1. For Fig. 1c, Extended Data Fig. 2b, Extended Data Fig. 2c, Extended Data Fig. 3, and Extended Data Fig. 6a the simulations for each parameter were performed 50 times, each with a different seed. The results were averaged over the 50 runs. Confounder regression was performed for the total number of transcripts per cell.

PBMC data^11^ was downloaded from the 10x genomics website (https://cf.10xgenomics.com/samples/cell/pbmc3k/pbmc3k_filtered_gene_bc_matrices.tar.gz). For the calculation of the tARI, clustering with Scanpy, TSCAN, *k*-means, and hierarchical clustering, preprocessing was performed with the *scanpy python package (V 1*.*4*.*6)* following the provided pipeline (https://scanpy-tutorials.readthedocs.io/en/latest/pbmc3k.html) for the filtering of cells and genes, normalization, and log-transformation as well as cluster annotation. For the clustering with Seurat, the provided Seurat pipeline was used (https://satijalab.org/seurat/archive/v3.2/pbmc3k_tutorial.html) for preprocessing, such as cell and gene filtering, normalization, log-transformation and cluster annotation using the Seurat

R package (*V 3*.*1*.*5*). CD8 T cells and B cells were extracted from the data and each cluster was standardized gene-wise and cell-wise before the calculation of the singular value decomposition. To remove any sub-structure in these clusters and before the reconstruction of the matrices from the SVD, singular values above the MP distribution were moved into the bulk, and the transcriptome mode (i.e. the singular vector that would have the largest singular value without normalization, see Supplementary Methods Note 1) was moved above the MP distribution. Then, two synthetic clusters containing 150 cells each were created from the cleaned-up original clusters. For cluster 1, a weighted average of a randomly picked B cell with expression profile *c*_*B*_ and a randomly picked CD8 T cell with expression profile *c*_*CD8T*_ was calculated according to: *c*_1_ = *α* · *c*_*B*_ + (1 − *α*) · *c*_*CD8T*_. For cluster 2, the weights were flipped: *c*_2_ = (1 − *α*) · *c*_*B*_ + *α* · *c*_*CD8T*_. *α* was chosen in a range from 0 to 1. *α* close to 0.5 produced highly similar clusters, while *α* close to 0 or 1 produced maximally different clusters (see Fig S2d). For each value of *α*, the procedure was repeated 50 times, each with a different seed for selecting 300 cells per cell type, and the results were averaged.

RNA-mix data^14^ was downloaded from the provided GitHub page. The data were normalized with the R scran package (V 1.14.6) and then log-transformed. Confounder regression was performed for the total number of transcripts, average mitochondrial gene expression, and average ribosomal gene expression. Two different merged clusters were created from the provided RNA mixtures as shown in Extended Data Fig. 5.

Bone marrow mononuclear cell data set (BMNC)^16^ was downloaded from the *R package SeuratData (bmcite, V 0*.*2*.*1)*. Normalization and the calculation of the G2M score^32^ were performed with the *Seurat R package (V 3*.*1*.*5*). Confounder regression was performed for the log-transformed total number of transcripts, cell cycle score, and average expressions of each: mitochondrial genes and ribosomal genes (list obtained from the HGNC website).

For the fetal kidney data set^19^, the same preprocessing and normalization was used as reported previously (scran R package^33^). The data was then log-transformed and the G2M score was calculated with the *Seurat R package*. Confounder regression was performed for the log-transformed total number of transcripts, G2M scores, and the average expressions of each: mitochondrial genes, ribosomal genes, and stress-related genes^34^.

### Embedding

Uniform Manifold Approximation and Projections^35^ (UMAPs) for individual clusters were calculated with the R package umap (V 0.2.7.0) on the first 10 PCs, 20 nearest neighbors, min_dist = 0.3, and Euclidean distances. The umap for BMNC data was calculated with the *Seurat R package* using 2000 highly variable genes (hvg), d = 50, k = 50, min.dist = 0.6 and metric = cosine. For the fetal kidney data set a force-directed graph layout was calculated using *the scanpy python package*. The graph was constructed using 100 nearest neighbors, 50 PCs, and the ForceAtlas2 layout for visualization.

### Differential expression test

Differentially expressed genes within the sub-clusters found in Extended Data Fig. 7 and Extended Data Fig. 8 were calculated with the function *findMarkers* of the *scran R package* on log-transformed normalized counts. Genes with a false discovery rate below 0.05 were selected and then sorted by log2 fold change. In Figures S7e and S8e, genes with the top 20 highest/lowest values in the gene singular vectors are listed and colored blue if they correspond to the top 20 DE genes.

### Staining

A human fetal kidney (female) at week 15 of gestation was used for immunofluorescence using the same procedure as reported previously^19^. The following primary antibodies were used: rabbit anti-UPK1A (1:35, HPA049879, Atlas Antibodies), mouse anti-KRT7 (1:200, #MA5-11986, Thermo Fisher Scientific), rabbit anti-CDH1 (1:50, SC-7870, Santa Cruz), rabbit anti-CLDN1 (1:100, #717800, Thermo Fisher Scientific), goat anti-CAV2 (1:100, AF5788-SP, R&D Systems), mouse anti-AKAP12 (1:50, sc-376740, Santa Cruz), rabbit anti-CLDN11 (1:50, HPA013166, Sigma Aldrich), mouse anti-POSTN (1:100, sc-398631, Santa Cruz) and goat anti-SULT1E1 (1:50, AF5545-SP, R&D Systems). The secondary antibodies were all purchased from Invitrogen and diluted to 1:500: Alexa Fluor 594 donkey anti-mouse (A21203), Alexa Fluor 594 donkey anti-rabbit (A21207), Alexa Fluor 647 donkey anti-mouse (A31571), Alexa Fluor 647 donkey anti-rabbit (A31573), Alexa Fluor 647 donkey anti-goat (A21447). The sections were imaged on a Nikon Ti-Eclipse epifluorescence microscope equipped with an Andor iXON Ultra 888 EMCCD camera (Nikon, Tokyo, Japan).

## Ethics statement

The collection and use of human material in this study was approved by the Medical Ethics Committee from the Leiden University Medical Center (P08.087). The gestational age was determined by ultrasonography, and the tissue was obtained from women undergoing elective abortion. The material was donated with written informed consent. Questions about the human material should be directed to S. M. Chuva de Sousa Lopes

## Data availability

The BMNC data can be downloaded with the R package SeuratData, named “bmcite”. The fetal kidney data is available with the SIGMA R package at https://github.com/Siliegia/SIGMA, named “sce_kidney”. The PBMC data can be downloaded at https://cf.10xgenomics.com/samples/cell/pbmc3k/pbmc3k_filtered_gene_bc_matrices.tar.gz and the RNA-mix data is available at https://github.com/LuyiTian/sc_mixology, named “mRNAmix_qc”.

## Code availability

The R package implementing SIGMA is available at https://github.com/Siliegia/SIGMA.

## Acknowledgments

We thank Ahmed Mahfouz for valuable feedback on the manuscript and the staff of Gynaikon, Rotterdam, as well as the anonymous tissue donors for the human fetal material.

## Author contributions

S.S., M.H., D.G., and M.M. developed the clusterability measure. M.M. designed and implemented the algorithms. S.S., M.H. and M.M. analyzed and interpreted the results.

S.M.S.L. provided the fetal kidney samples. X.F. sectioned and performed the immunostaining of the fetal kidney samples. M.H. imaged the kidney samples and interpreted the imaging results. S.S., M.H. and M.M. wrote the manuscript with contributions from D.G. All authors have read and approved the final version of the manuscript.

## Competing Interests statement

The authors declare no competing interests.

## Funding information

M.M, S.S. and D.G. were supported by the Netherlands Organisation for Scientific Research (NWO/OCW, www.nwo.nl), as part of the Frontiers of Nanoscience (NanoFront) program. We acknowledge funding by an NWO/OCW Vidi grant (016.Vidi.189.007) for M.H. and S.S., by China Scholarship Council (CSC 201706320328) to X.F and by a European Research Council Consolidator Grant OVOGROWTH (ERC-CoG-2016-725722) to S.C.S.L. This work was carried out on the Dutch national infrastructure with the support of SURF Foundation.

## Figures

**Extended Data Fig. 1.**
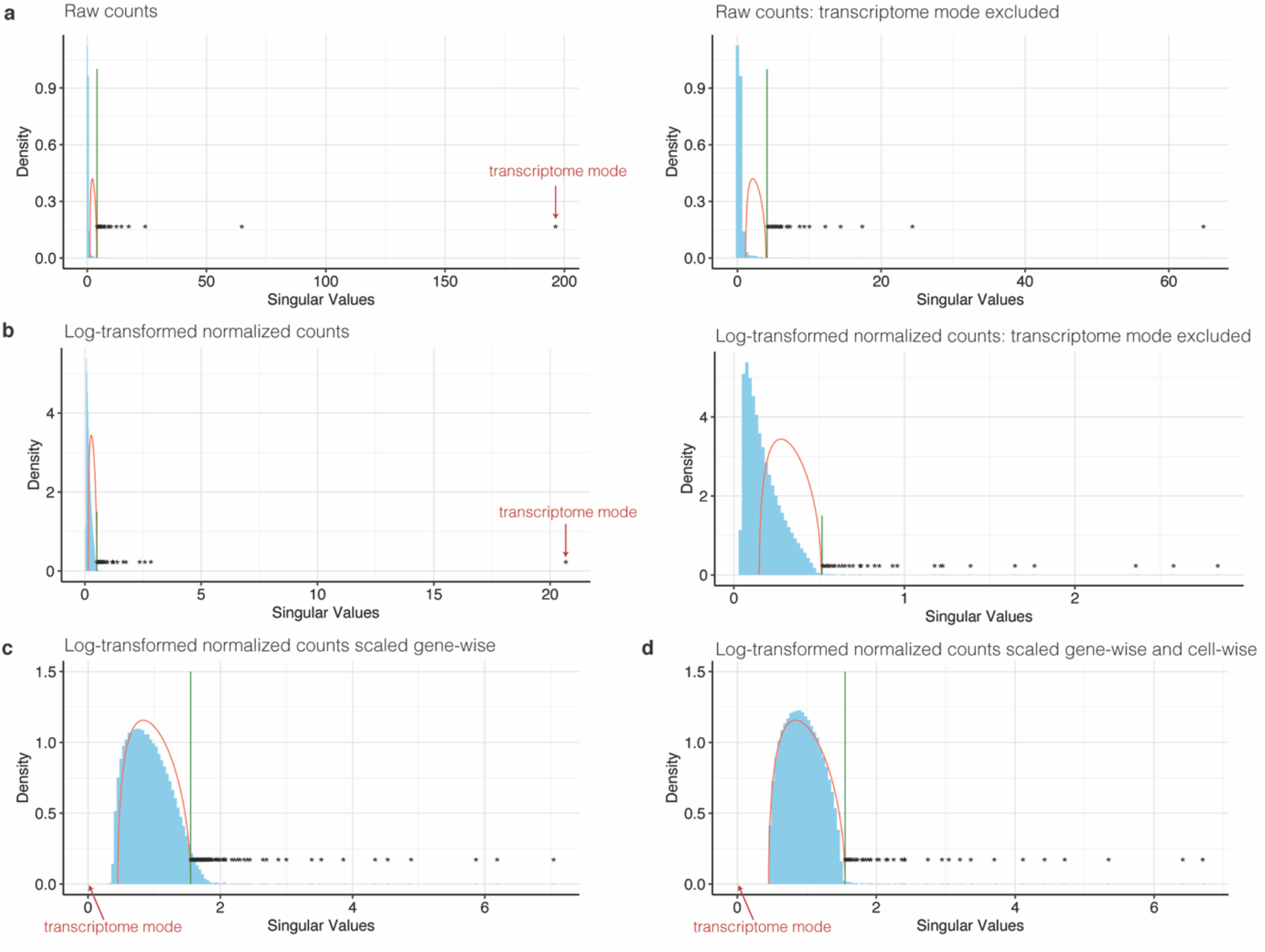
Random matrix theory can be applied to measured single-cell RNA-seq data sets after proper pre-processing. SV spectra of the fetal kidney single-cell RNA-seq data set after different preprocessing steps. **a** Raw UMI counts. Arrow indicates transcriptome mode. Right: Transcriptome mode was excluded. The bulk of the SV spectrum does not coincide with the MP distribution. **b** Log-transformed, normalized UMI counts. Arrow indicates transcriptome mode. Right: Transcriptome mode was excluded. The SV spectrum does not coincide with the MP distribution. **c** Log-transformed, normalized data as in b, that were additionally centered gene-wise. The SV spectrum approximately coincides with the bulk of the MP distribution and the transcriptome mode, visible as the highest singular value in b and c appears close to 0 (indicated by the arrow). **d** Log-transformed, normalized, and gene-wise standardized data, as in c, that was additionally standardized cell-wise. The SV spectrum coincides with the bulk of the MP distribution. There are no free parameters to fit. The MP distribution is fully determined by the number of measured genes and cells.

**Extended Data Fig. 2.**
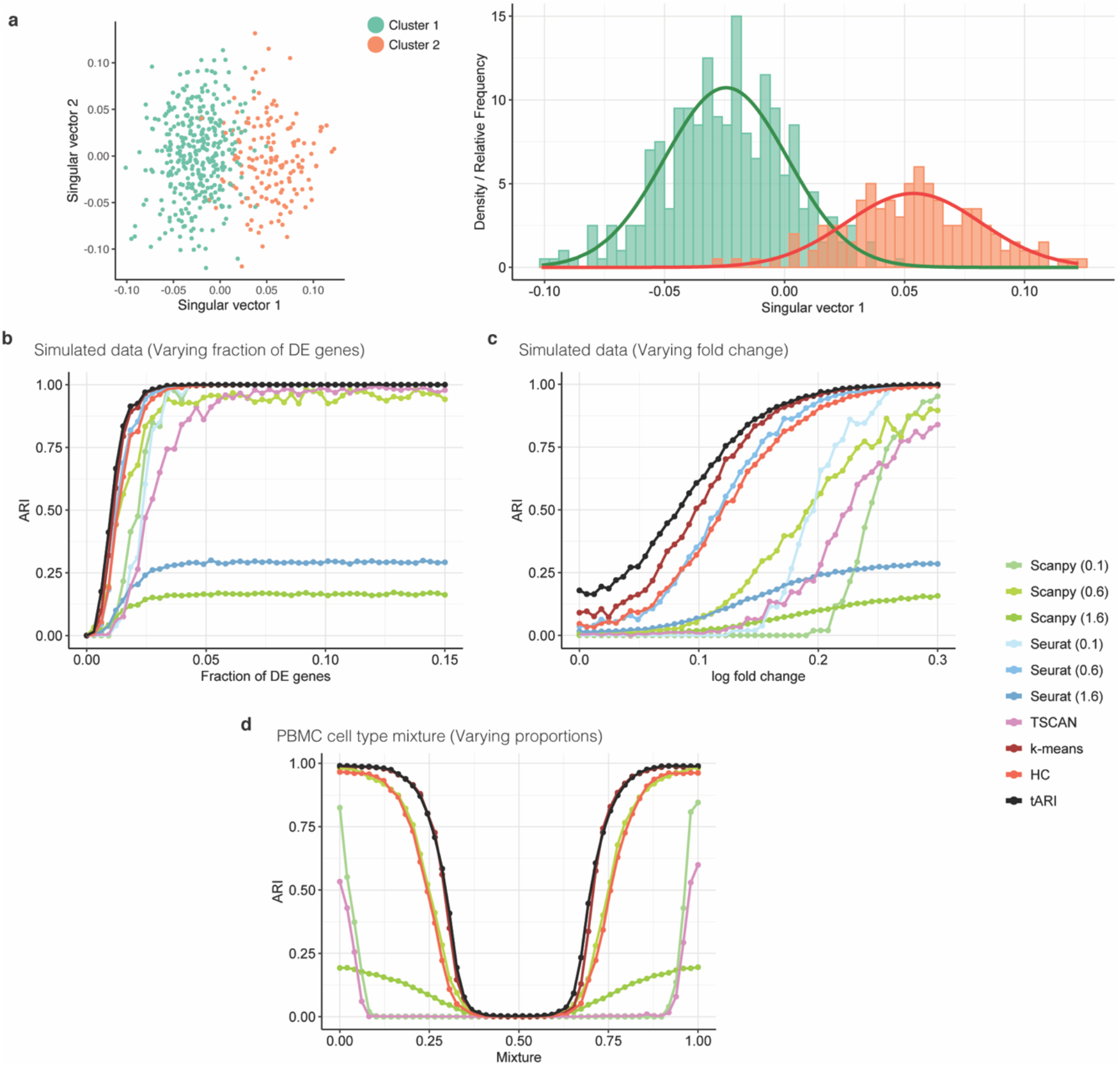
An approximate upper limit to the achievable ARI can be derived from an estimate of the Bayesian error rate. **a** First two singular vectors of a simulated data set with two clusters. Only the first singular vector is significant. Right: Histogram of the first singular vector. The color indicates to which simulated (ground truth) cluster the cells belong. Two normal distributions fitted separately to the singular vector entries belonging to the two clusters are shown as solid lines. The Bayesian error rate is estimated from the overlap of these two distributions and used to calculate the theoretical ARI (tARI). **b** ARI achieved by various clustering methods compared to the ground truth and tARI for simulated data with two clusters. The number of differentially expressed genes was varied. **c** ARI achieved by various clustering methods compared to the ground truth and tARI for simulated data with two clusters. The log fold change between clusters was varied. **d** ARI achieved by various clustering methods compared to the ground truth and tARI for PBMC cell-type mixture. The mixture proportions were varied from 0 to 1. **b**,**c**,**d** The numbers in the legend indicate the resolution parameter used.

**Extended Data Fig. 3.**
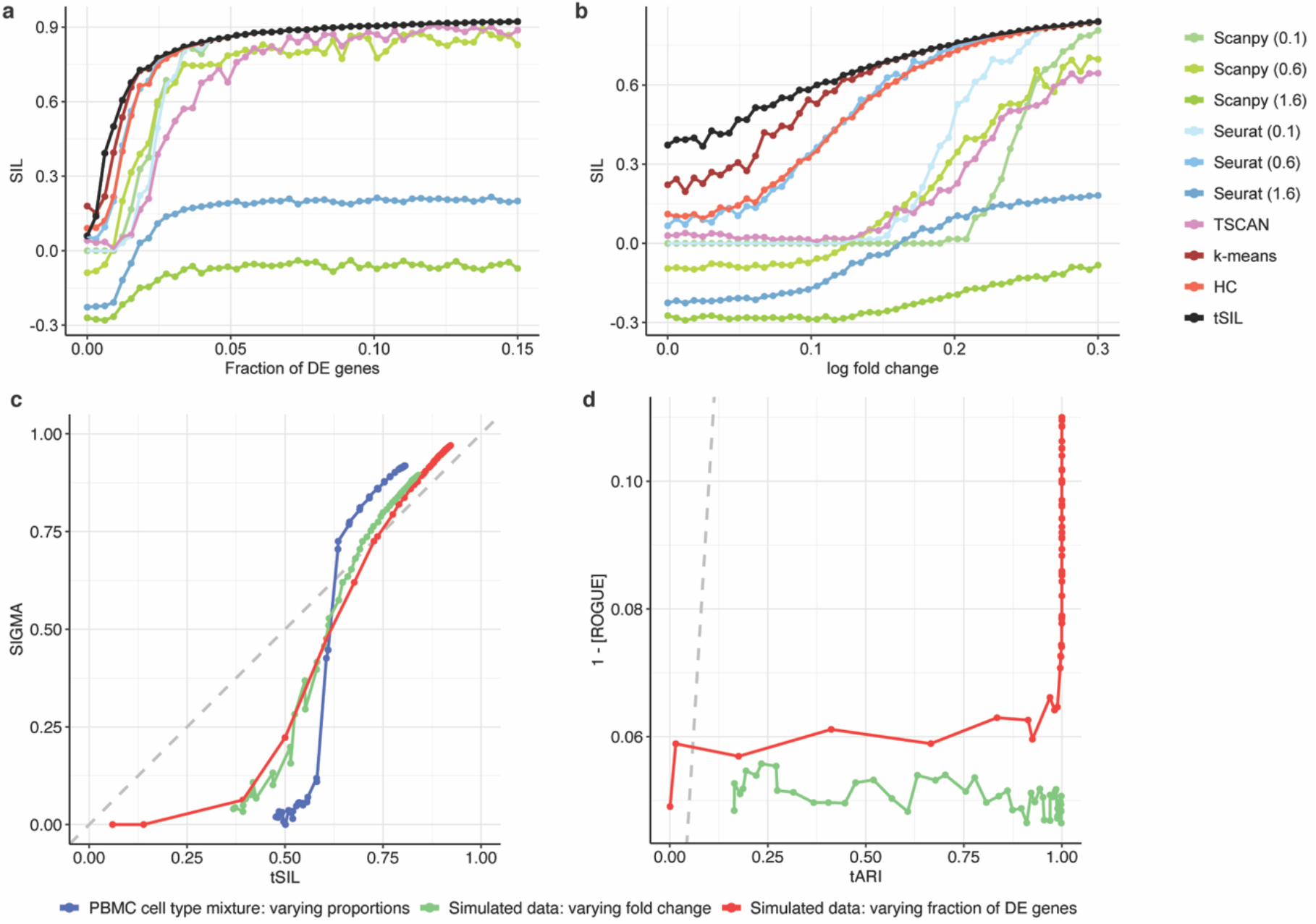
An approximate upper limit to the best possible silhouette coefficient and accordance of ROGUE with tARI. **a** Silhouette coefficient (SIL) achieved by various clustering methods and theoretical SIL (tSIL) for simulated data with two clusters. The number of differentially expressed (DE) genes was varied. **b** SIL achieved by various clustering methods and tSIL for simulated data with two clusters. The log fold change between clusters was varied. **a**,**b** The numbers in the legend indicate the resolution parameter used. **c** tSIL versus SIGMA. Red data points: Simulated data sets with two clusters. The number of DE genes was varied, the log fold change between clusters was fixed. Green data points: Simulated data sets with two clusters. The log fold change between clusters was varied, the number of DE genes was fixed. Blue data points: Two synthetic clusters were created by weighted averages of cells from two clusters in the PBMC data set (see Fig. 2c). Cluster weights were varied. The Grey dashed line indicates identity. **d** tARI versus 1 - [ROGUE] score. Red data points: Simulated data sets with two clusters. The number of DE genes was varied, the log fold change between clusters was fixed. Green data points: Simulated data sets with two clusters. The log fold change between clusters was varied, the number of DE genes was fixed. Blue data points: Two synthetic clusters were created by weighted averages of cells from two clusters in a PBMC data set (see Fig. 2c). Cluster weights were varied. The Grey dashed line indicates identity.

**Extended Data Fig. 4.**
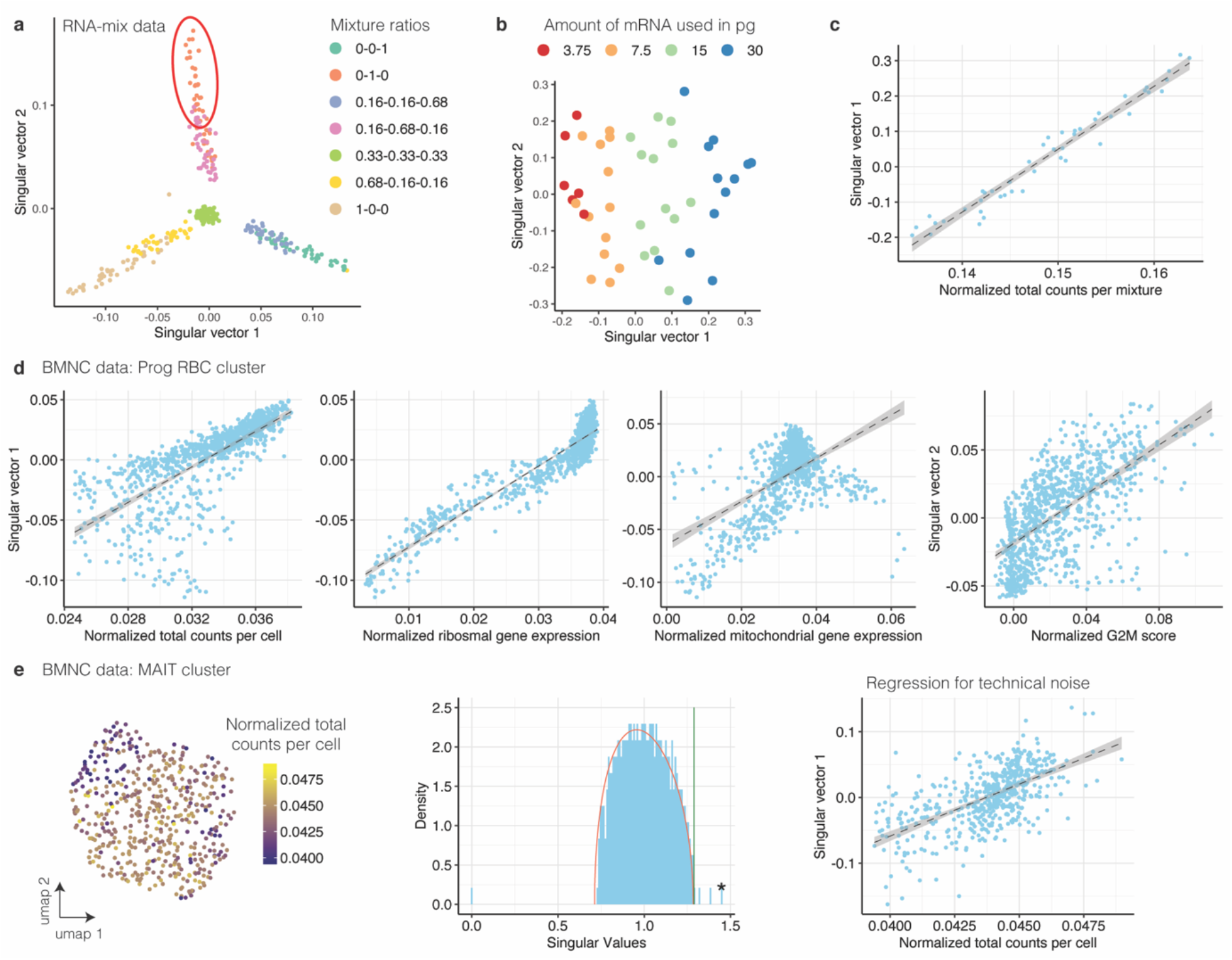
Singular vectors from measured data contain confounding technical variability. The singular vectors are correlated with several confounding variables. **a** First two singular vectors of the RNA-mix data set. Clusters are indicated by color. **b** First two singular vectors of the cluster indicated by a red solid ellipse in a. The amount of mRNA per mixture [pg] is indicated in color. **c** Normalized total counts per mixture versus first singular vector of the cluster shown in b. Linear regression (dashed line) is used to regress out the correlation with the total counts. Grey area indicates standard deviation. **d** First singular vector of Prog RBC cluster in the BMNC data set versus normalized total counts per cell, normalized expression of ribosomal genes, and normalized expression of mitochondrial genes. Right: Second singular vector versus normalized G2M score. The dashed line indicates the linear regression and the grey area indicates the standard deviation. **e** Left: UMAP of MAIT cluster in BMNC data set. The color indicates the normalized total counts per cell. Middle: SV spectrum and MP distribution of the MAIT cluster. Only 1 significant singular value is indicated by an asterisk. Right: Normalized total counts per cell versus the singular vector associated with the significant singular value (here: 1st singular vector) in the MAIT cluster.

**Extended Data Fig. 5.**
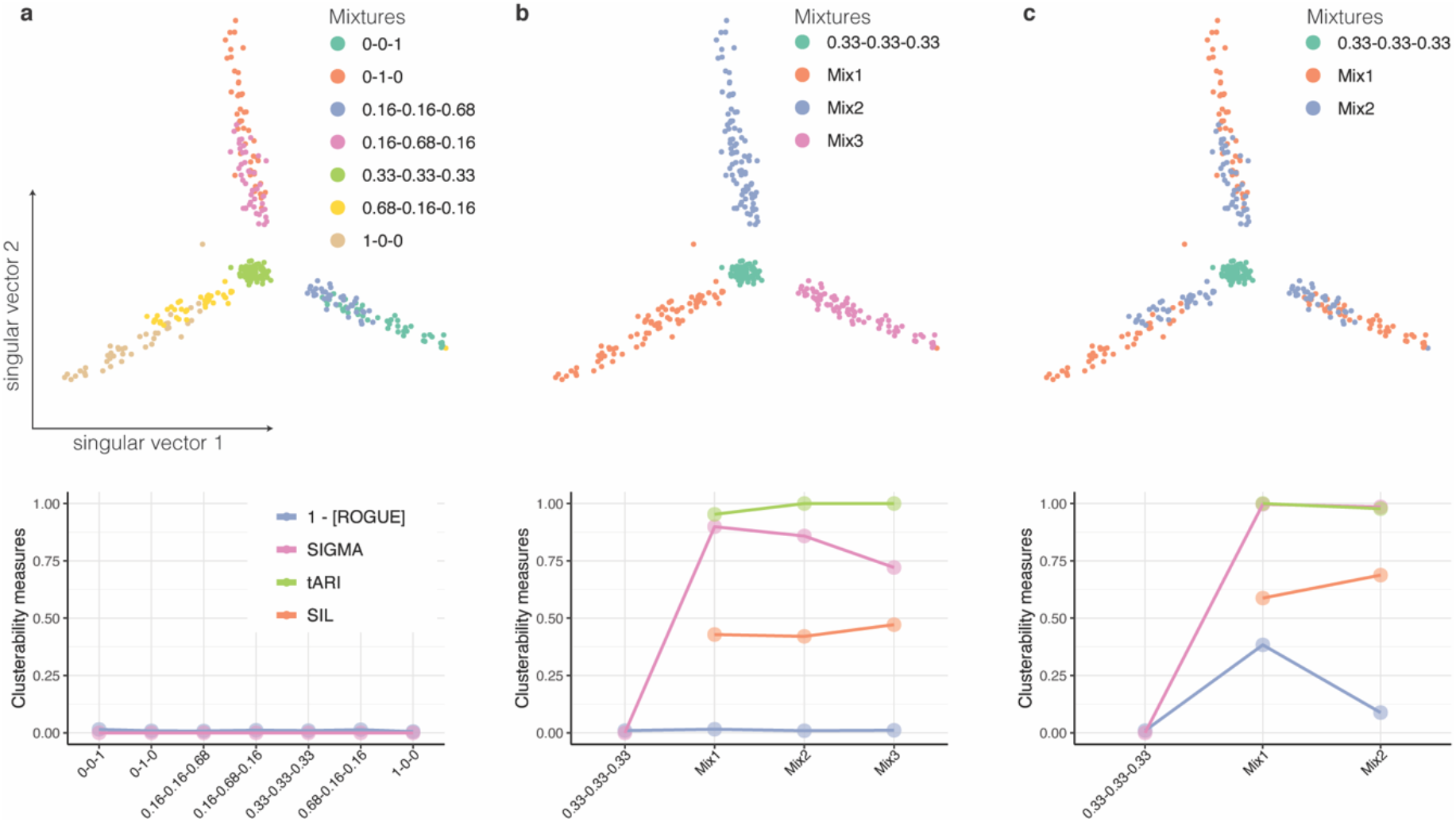
SIGMA outperforms other measures on experimental data. Clusters were merged in different ways to vary the amount of variability in each merged cluster. Top: first two singular vectors of RNA-mix data. Colors indicate different mixtures. Bottom: The values of SIGMA (rose), SIL (orange), tARI (green) and 1 - [ROGUE] (blue) for each corresponding cluster. **a** Original RNA mixture. **b** Merged clusters. Blue: 0-1-0 merged with 0.16-0.68-0.16. Pink: 0-0-1 merged with 0.16-0.16-0.68. Orange: 1-0-0 merged with 0.68-0.16-0.16. **c** Blue merged cluster contains mixtures 0.68-0.16-0.16, 0.16-0.68-0.16 and 0.16-0.16-0.68. Orange merged cluster contains mixtures 1-0-0, 0-1-0, and 0-0-1.

**Extended Data Fig. 6.**
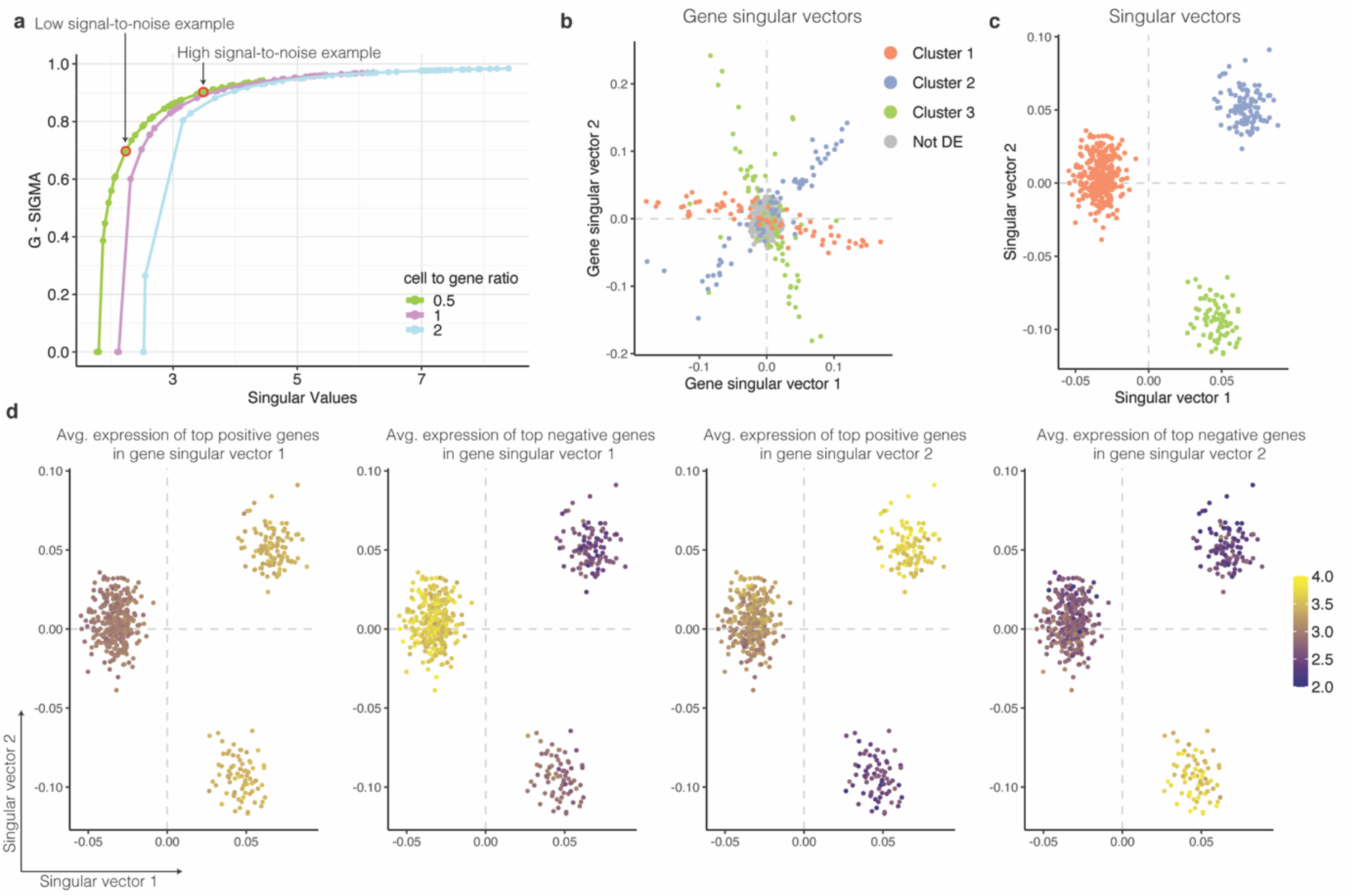
Variance-driving genes identified by random matrix theory coincide with differentially expressed genes in a simulated data set. Genes with high absolute values in the gene singular vector contribute the most to the variability. **a** Value of the largest singular value versus the squared cosine of the angle between the gene singular vector of the signal matrix and the gene singular vector of the measured expression matrix (G-SIGMA) in simulated data. Arrows indicate examples shown in Figure 2a. **b** First two gene singular vectors. Differentially expressed genes of each cluster are indicated by color. **c** First two singular vectors for the simulated data set shown in panel b. Dashed grey lines indicate the 0 value on each of the axes. Cell clusters are indicated by color. **d** First two singular vectors as in c. Dashed grey lines indicate the 0 value on each of the axes. The average log-transformed expression of the top 1% genes driving the variance is indicated by color. The 4 panels show, respectively, from left to right: genes corresponding to the highest values in gene singular vector 1, genes corresponding to the lowest values in gene singular vector 1, genes corresponding to the highest values in gene singular vector 2, and genes corresponding to the lowest values in gene singular vector 2.

**Extended Data Fig. 7.**
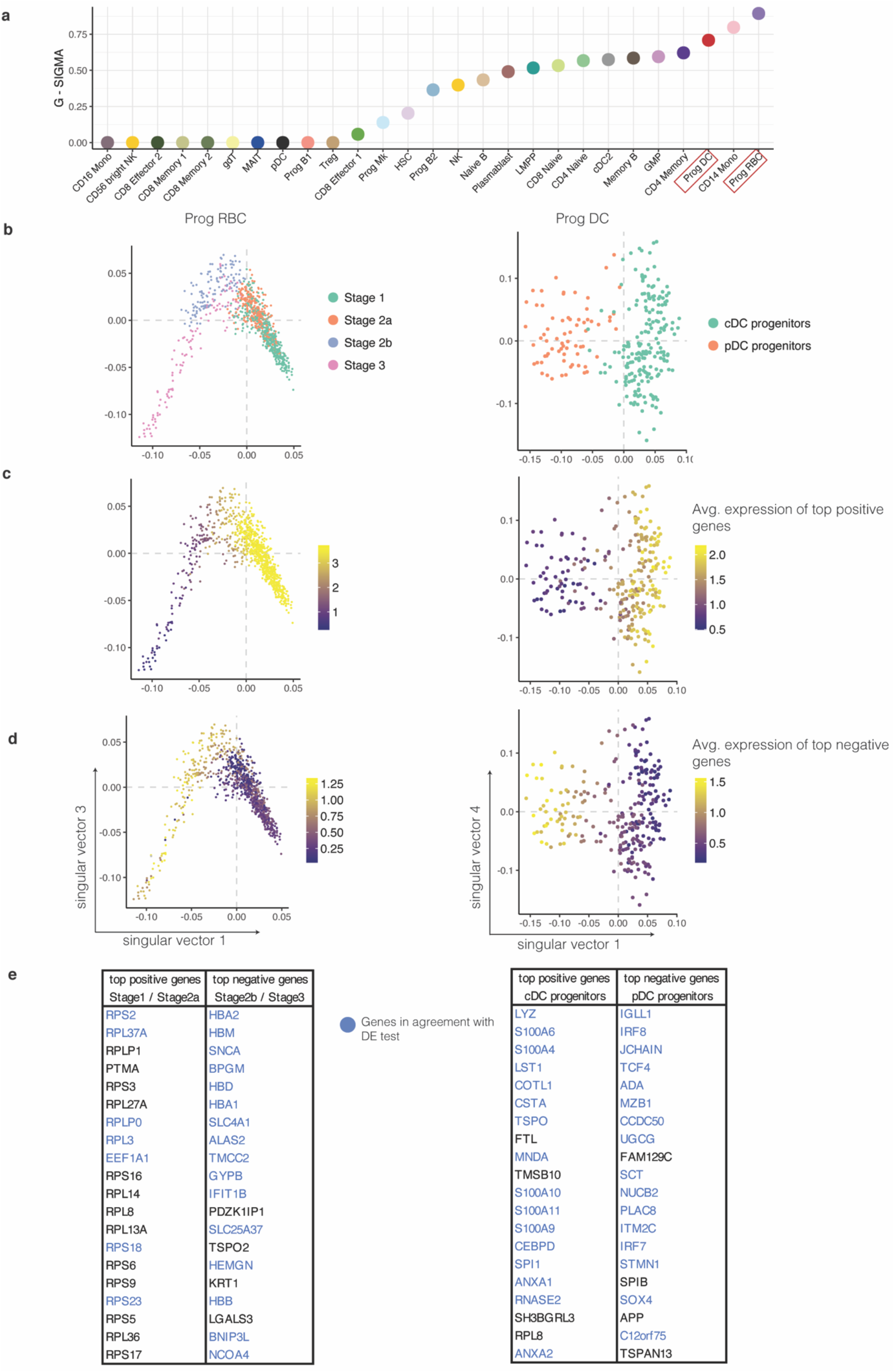
Congruence between variance-driving genes and differentially expressed genes between sub-clusters in the fetal kidney. **a** G-SIGMA for each cluster in the BMNC data set. **b** Singular vectors of the two clusters from the BMNC data set with the highest SIGMA. The color indicates sub-clustering. Dashed grey lines indicate the 0 value on each of the axes. **c** Singular vectors of clusters shown in panel a with color indicating the average log-transformed gene expression of genes with the 1% highest values in the first gene singular vector. **d** Singular vectors of clusters shown in panel a with color indicating the average log-transformed gene expression of genes with the 1% lowest values in the first gene singular vector. **e** Genes driving the variance in the two clusters shown in b. These genes have the 20 highest/lowest values in the first gene singular vector respectively. In blue: top 20 upregulated genes based on differential expression (DE) test between the sub-clusters using *findMarkers* (from *scran* R package).

**Extended Data Fig. 8.**
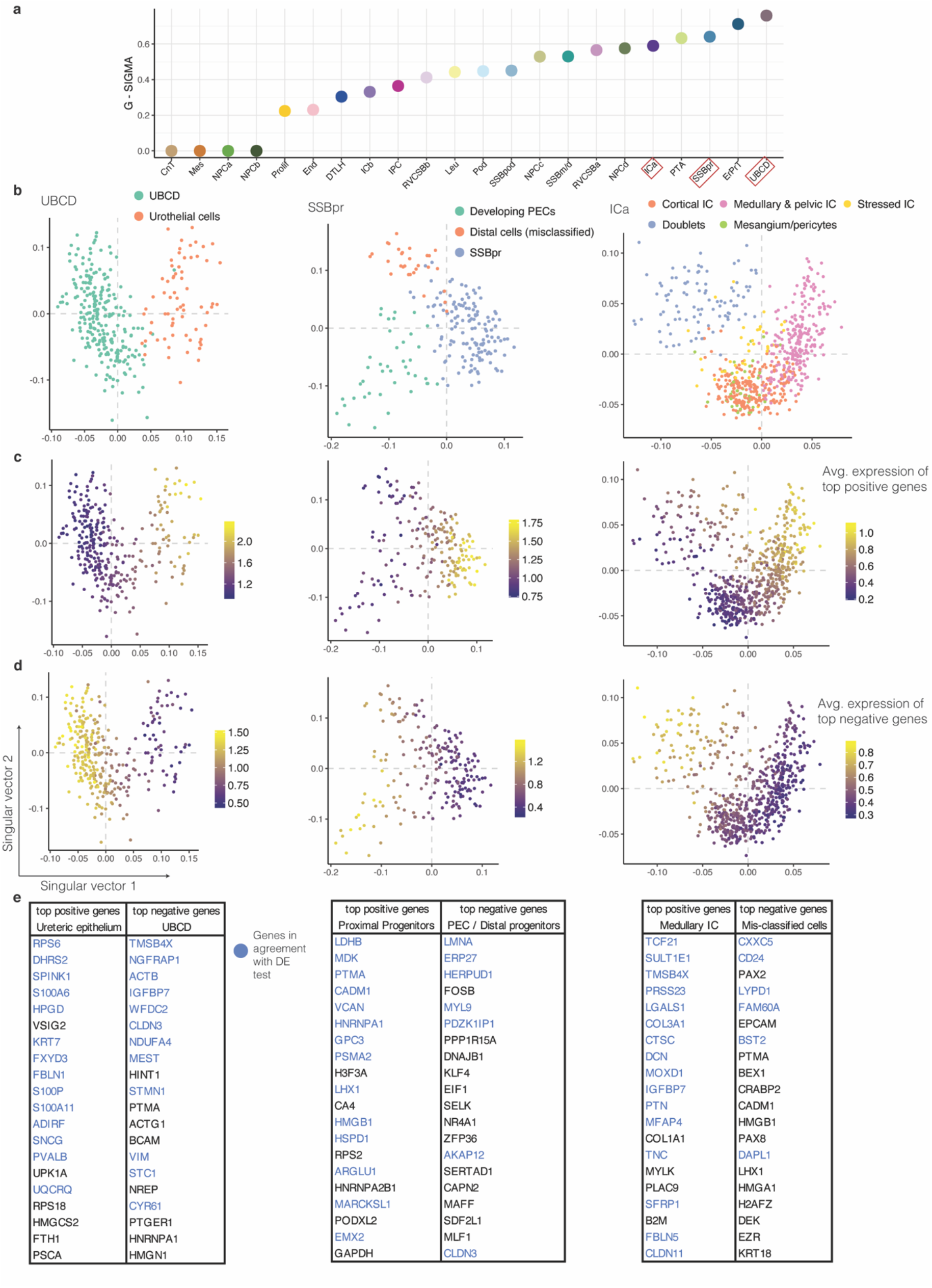
SIGMA indicates sub-structures in a fetal kidney data set. **a** G-SIGMA for each cluster in the fetal kidney data set. **b** First two singular vectors of the three clusters from the fetal kidney data set with high SIGMA. The color indicates sub-clustering. Dashed grey lines indicate the 0 value on each of the axes. **c** First two singular vectors of clusters shown in panel a with color indicating the average log-transformed gene expression of genes with the 1% highest values in the first gene singular vector. **d** First two singular vectors of clusters shown in panel a with color indicating the average log-transformed gene expression of genes with the 1% lowest values in the first gene singular vector. **e** Genes driving the variance in the three clusters shown in b. These genes have the 20 highest/lowest values in the first gene singular vector respectively. In blue: top 20 upregulated genes based on differential expression (DE) test between the sub-clusters using *findMarkers* (from *scran* R package).

**Extended Data Fig. 9.**
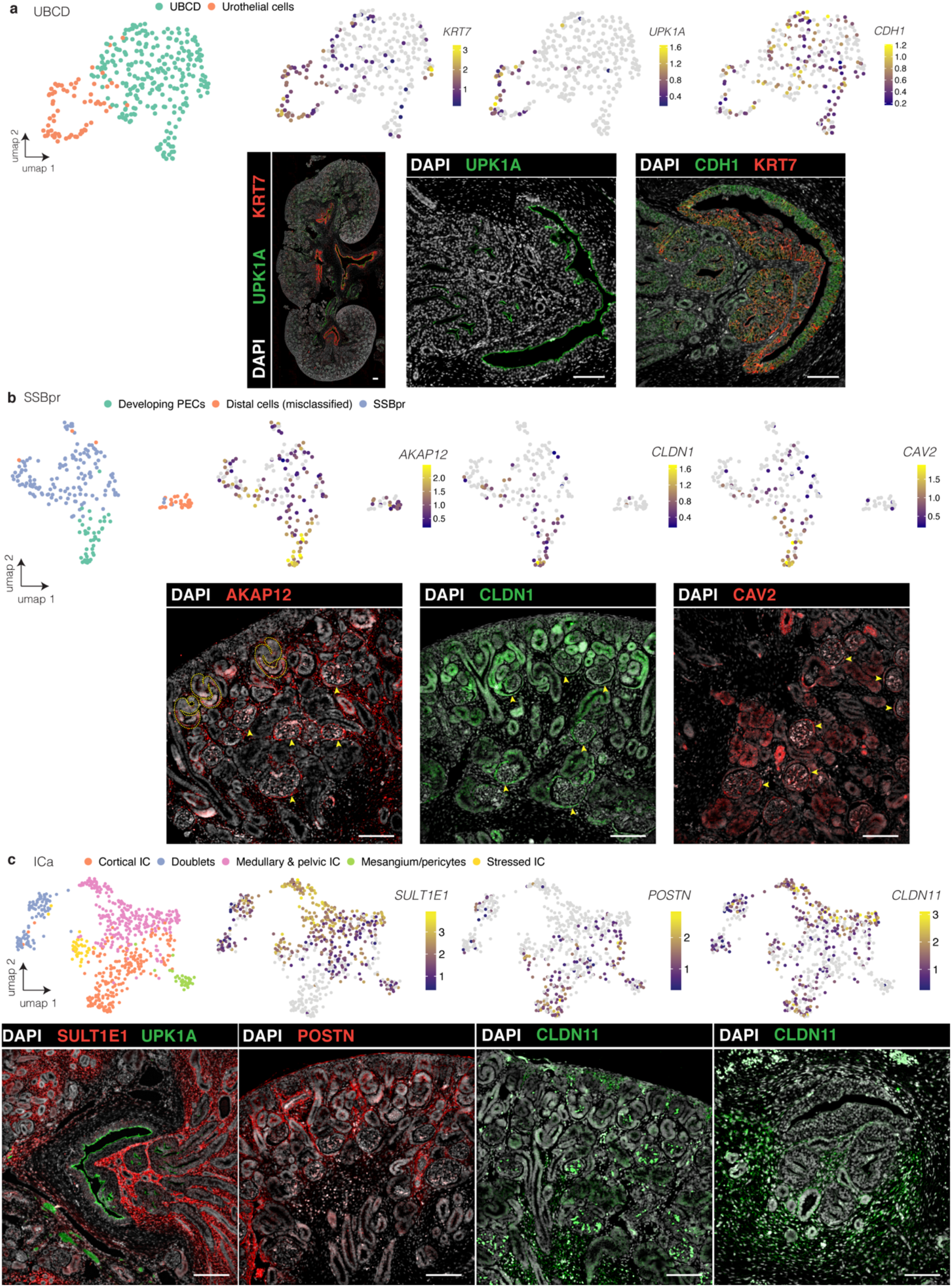
Immunostaining validates newly identified subclusters in fetal kidney data set. **a-c** Upper panels show UMAPs of the selected clusters in the fetal kidney data set. Log-normalized expression of selected genes is indicated by color. Lower panels show immunostainings of week 15 fetal kidney sections. **a** UBCD cluster. UPK1A, CDH1, and KRT7 expressions are shown in a complete section (leftmost image) and in the urothelial epithelium. **b** SSBpr cluster. Expression of AKAP12, CLDN1 and CAV2 is shown. The dashed lines indicate S-shaped bodies, arrows indicate PECs in developing glomeruli **c** ICa cluster. Expression of SULT1E1 and UPK1A is shown around the ureter expression of POSTN is shown in cortical areas, CLDN11 is shown in the cortical area (CLDN11, left image) and around the ureter (CLDN11, right image). Scale bars: 100 µm.

## Notes

### Competing Interest Statement

The authors have declared no competing interest.

