## Supplementary Information for "A clusterability measure for single-cell transcriptomics reveals phenotypic subpopulations"

### Supplementary Note 1

**4 References** **12**

#### 1 Introduction

Our aim is to develop a clusterability measure for scRNA-seq data. As we define more precisely in section (2.5), we consider clusterability to be the clustering quality that is optimally achievable, given a certain amount of noise in the data. Clustering quality can only be assessed quantitatively if the ground truth is known, which is strictly only the case for simulated data. A clusterability measure must thus be able to reflect clustering quality without knowledge of the ground truth. Such a measure would be highly useful, since it would allow us to detect the presence of meaningful (non-random) variability, and thus determine the necessity to sub-cluster measured data. For the development of this clusterability measure we will use concepts from random matrix theory and perturbation theory. In short, we decompose the single-cell gene expression matrix $\tilde{X}$ into a random matrix $X$, which contains technical and biological noise, and a signal matrix $P$, which contains the expression profiles of different cell types or states. Then, we apply perturbation theory, treating the signal matrix $P$ as a low-rank perturbation of the noise matrix $X$. Perturbation theory then allows us to calculate the angle between the singular vectors of the measured single cell expression matrix $\tilde{X}$ and the corresponding singular vectors of the unobserved signal matrix $P$. The cosine of this angle constitutes a useful clusterability measure because a large value (small angle) indicates a high signal-to-noise ratio (and thus high clusterability) and a small value (large angle) indicates a low signal-to-noise-ratio (and thus low clusterability). We show empirically that this clusterability measure is a proxy for the

theoretically achievable adjusted rand index [Figure 1d].
In what follows, we first present our model of gene expression data (2.1) and introduce matrix decomposition (2.2). Subsequently, we introduce the Marchenko-Pastur (MP) distribution (2.3), which describes the eigenvalue spectrum of a random matrix and apply perturbation theory to link the (unobserved) signal matrix to the spectrum of the measured expression matrix (2.4). In section (2.5), we establish our notion of clusterability. In section (3.1), we describe the preprocessing steps necessary for the application of the theory to single-cell RNAseq data. Then, in section (3.2), we develop a method to remove the effect of nuisance variables (i.e. sources of systematic, non-random variability that should not drive clustering.) The complete algorithm can be found in section (3.3).

### 2 Signal-Measurement-Angle (SIGMA)

#### 2.1 Model

Let $\tilde{X} \in \mathbb{R}^{M\times N}$ be the measured single-cell expression matrix with $M$ the number of genes (rows) and $N$ the number of cells (columns). We model the measurement $\tilde{X}$ as the sum of a random noise matrix $X \in \mathbb{R}^{M\times N}$ and a "signal" matrix $P \in \mathbb{R}^{M\times N}$.

$$\tilde{X} = X + P \tag{1}$$

In our model, $X$ contains both technical and biological noise. For example, if there was only one cell type or cell state present in a data set, $P$ would consist of identical columns. Note that we only observe the matrix $\tilde{X}$ experimentally. We will show below, that we can make a statement about the influence of the noise $X$ on the signal $P$, without knowing $X$ or $P$. To achieve that we invert the logic of conventional models: instead of modeling the influence of random noise on the signal, we consider the influence of a deterministic perturbation on a random matrix. All results rely on matrix decomposition, which will be introduced next.

#### 2.2 Matrix decomposition

##### 2.2.1 Eigendecomposition

We first define the cell-cell correlation matrix. To that end, we assume that $\tilde{X}$ has been standardized cell-wise (i.e. column-wise) to mean 0 and standard deviation 1. The cell-cell correlation matrix $C \in [-1, 1]^{N\times N}$ is then defined as:

$$C = \frac{1}{M-1}\tilde{X}^T\tilde{X} \tag{2}$$

The correlation matrix is a square and symmetric matrix which can hence, by the spectral theorem, undergo eigendecomposition into the form

$$C = V\Sigma V^T = \sum_{i=1}^{N} \lambda_i v_i v_i^T. \tag{3}$$

$V \in \mathbb{R}^{N\times N}$ contains the eigenvectors $v_i$ of $C$ in the columns and $\Sigma \in \mathbb{R}^{N\times N}$ is a diagonal matrix containing the eigenvalues $\lambda_i$ of $C$. If $M < N$, then $C$ is a singular matrix and will contain at least $N - M$ eigenvalues equal to 0, which is an important consideration for the definition of the Marchenko-Pastur distribution (see below).
In full analogy to the cell-cell correlation matrix we can define a gene-gene correlation matrix $\hat{C}$, now assuming that the expression matrix $\tilde{X}$ has been standardized gene-wise (row-wise) to mean 0 and standard deviation 1:

$$\hat{C} = \frac{1}{N-1}\tilde{X}\tilde{X}^T. \tag{4}$$

If $M > N$, then $\hat{C}$ is a singular matrix and will contain at least $M - N$ eigenvalues equal to 0. Therefore either $C$ (if $M < N$) or $\hat{C}$ (if $M > N$) is a singular matrix (unless $M = N$) with at least $|N - M|$ eigenvalues equal to 0.

##### 2.2.2 Singular value decomposition

To decompose the (rectangular) expression matrix $\tilde{X}$ into noise and signal, we use singular value decomposition:

$$\tilde{X} = \sum_{i=1}^{N} \gamma_i u_i v_i^T.$$

The $v_i$'s are the right singular vectors of $\tilde{X}$ and correspond to the eigenvectors of the cell-cell correlation matrix. We will call them singular vectors in the following. The $u_i$'s are the left singular vectors of $\tilde{X}$ and correspond to the eigenvectors of the gene-gene correlation matrix, which we will call gene singular vectors. The singular values are denoted by $\gamma_i$. The singular values of $\tilde{X}$ and the eigenvalues of the corresponding correlation matrix have a known connection given by:

$$\lambda_i = \gamma_i^2.$$

#### 2.3 Random Matrix Theory

The Marchenko-Pastur (MP) distribution is widely used to reveal nonrandom properties of empirical correlation matrices in physics and finance [1, 2]. The MP distribution describes the distribution of eigenvalues of a random correlation matrix in the asymptotic limit [3, 4, 5] (for $N \to \infty$ and $M \to \infty$, $\frac{N}{M} < 1$). The entries of the random matrix are arbitrary as long as they are distributed identically and independently. scRNA-seq data are typically modeled by a Poisson, a negative binomial or a zero-inflated negative binomial distribution, which are in principle admissible in random matrix theory.

**Theorem 1 (Marchenko-Pastur)** *([3, 4, 5]) Let $Y$ be a $M \times N$ matrix with entries that are independent identically distributed (i.i.d.), mean 0 and variance $\nu^2 < \infty$. The corresponding Wishart matrix is defined as $W = \frac{1}{M} Y^T Y$. For $N \to \infty$, $M \to \infty$ and $0 < c < 1$, where $c$ is defined as $\frac{N}{M}$. The distribution of the eigenvalues $\lambda$ of $W$ is given by*

$$\mu(\lambda) = \frac{\sqrt{(b-\lambda)(\lambda-a)}}{2\pi c \lambda \nu^2}\, d\lambda \quad \text{if} \quad a \leq \lambda \leq b$$

*For $c > 1$ the distribution has an additional number of 0 eigenvalues:*

$$\mu(\lambda) = \frac{\sqrt{(b-\lambda)(\lambda-a)}}{2\pi c \lambda \nu^2} \mathbb{1}_{[a,b]} + (1 - \frac{1}{c})\delta_0(\lambda)$$

*with*

$$a, b = \nu^2 \left[1 \pm \sqrt{c}\right]^2.$$

$\delta_0(\lambda)$ is the Dirac delta function, which is 1 if $\lambda = 0$ and 0 otherwise. For the correlation matrix we obtain $\nu = 1$ because the mean of all eigenvalues is 1.
This theorem places the eigenvalues of a random correlation matrix into a compact interval between $[a, b]$. All eigenvalues of an empirical correlation matrix that fall within this interval can be considered to be due to random noise. The presence of eigenvalues above this distribution

indicates the existence of non-random structure in the data. An empirical (measured) correlation matrix can therefore be decomposed into a random part $C^r$ and a signal part $C^s$[3]:

$$C = \sum_{\lambda \leq b} \lambda_i v_i v_i^T + \sum_{\lambda > b} \lambda_i v_i v_i^T = C^r + C^s$$

$C^s$ contains the non-random and therefore biologically relevant correlations.

For the application of the MP distribution to an empirical correlation matrix we need to consider that the eigenvalues of a correlation matrix always sum up to 1. Thus, if there are eigenvalues above the MP distribution the bulk of the distribution (which is described by MP) will shift to the left. To approximately account for this shift, we introduce a modified MP-distribution as follows:

$$\mu^*(\lambda) = \frac{\mu(\lambda)}{\alpha},$$
$$a^* = \alpha a, \;\; b^* = \alpha b.$$

where $\alpha = 1 - \frac{\lambda_{max}}{N}$ and $a^*$ and $b^*$ replace $a$ and $b$ respectively.

We can formulate the MP distribution also for singular values, via a variable transform, and obtain the following density:

$$d\rho(\gamma) = \frac{\sqrt{(b-\gamma^2)(\gamma^2 - a)}}{\pi \gamma c}\, d\gamma \;\; \text{if} \;\; \sqrt{a} \leq \gamma \leq \sqrt{b} \tag{5}$$

In this case, all singular values that lie within the compact interval of $[\sqrt{a}, \sqrt{b}]$ can be considered to arise from random noise and singular values above this threshold indicate deterministic biological relevant signal. Thus, we can decompose the matrix $\tilde{X}$ into two parts:

$$\tilde{X} = \sum_{\gamma \leq \sqrt{b}} \gamma_i u_i v_i^T + \sum_{\gamma > \sqrt{b}} \gamma_i u_i v_i^T = \tilde{X}^r + \tilde{X}^s \tag{6}$$

The first part $\tilde{X}^r$ is random noise, the second part $\tilde{X}^s$ contains relevant signal.

The MP theorem holds strictly only in the asymptotic limit, but provides a very good approximation for big enough $N$ and $M$. For finite dimensions, there is however a non-zero probability that a random i.i.d matrix has eigenvalues above the MP distribution. That probability is described by the Tracy-Widom (TW) distribution.

**Theorem 2 (Tracy-Widom)** *([4, 6]) For empirical correlation matrices of size $N \times N$ of i.i.d. random variables with a finite fourth moment, the distance between the upper edge of the spectrum of the MP distribution $b$ and the largest eigenvalue $\lambda_{max}$ converges towards the Tracy-Widom distribution*

$$\text{Prob}\left(\lambda_{\max} \leq b + \gamma N^{-2/3} u\right) = F_1(u),$$

*where $\gamma$ in this case is given by $\gamma = \sqrt{c}\; b^{2/3}$.*

$F_1(u)$ is the TW distribution, the probability distribution of the re-scaled eigenvalues of a random Hermitian matrix. We are interested in the type-1 distribution which holds for Gaussian orthogonal ensembles [7]. The distribution function can not be explicitly stated but relies on numerical approximations.

The TW distribution can be formulated, as well, for the singular values via the variable transform:

$$\text{Prob}\left(\gamma_{max} \leq \sqrt{b + \gamma N^{-2/3} u}\right) = F_1(u), \tag{7}$$

Since we always work with finite matrices in practice, we use the TW distribution to discriminate between singular values that belong to noise and signal, respectively. Specifically, we use $u = 1$ as a cutoff, so that $F_1(1) \approx 0.95$. In other words, there is a probability of 0.05 that a singular value bigger than $\sqrt{b + \gamma N^{-2/3}}$ is observed, if the matrix is entirely random. If $N$ is very low, the MP distribution is not a good approximation anymore. For $N < 50$, we create an empirical distribution of noise-related singular values, by permuting the entries of the measured expression matrix $\tilde{X}$. For each permutation we calculate the singular values and note the largest singular value. The 95th quantile of the distribution of the largest singular values across permutations is then taken to be the cutoff between singular values stemming from noise and signal respectively.

To discriminate random from non-random matrix components we can also look at the singular vectors [8]. Singular vectors that correspond to random components are "de-localized" and their elements have the following distribution:

$$f(\psi) = (1 - \psi^2)^{\frac{N-3}{2}}$$

If $N$ is large, this distribution can be estimated by a Gaussian distribution with mean zero and variance $\frac{1}{N}$.

$$f(\psi) \sim \frac{N}{\sqrt{2\pi}} e^{\frac{-N\psi^2}{2}} \tag{8}$$

In order to distinguish localized from de-localized singular vectors, we can therefore assess the normality of the singular vectors. In our implementation we use a Shapiro-Wilk test. We assign singular vectors that obtain a p-value $< 0.01$ or are associated to singular values far from the bulk (the highest 50% of signal singular values) to real variability above the MP distribution.

#### 2.4 Perturbation theory

As explained above, we model the observed expression matrix $\tilde{X}$ as a random matrix $X$ perturbed by a deterministic signal matrix $P$. There is an important difference between the perturbation matrix $P$ in equation 1 and the matrix $\tilde{X}^s$ in equation 6. $\tilde{X}^s$ does contain biologically relevant information, but is still influenced by the effects of random noise, whereas the matrix $P$ consists of the pure signal without any added noise. The only case where these two matrices are identical is when the noise matrix $X$ and the perturbation matrix $P$ are spanned by the same singular vectors, which is rarely the case. It is thus not possible to recover the unobserved, noise-free signal matrix by using those singular vectors that are associated with the highest singular values.

While it is not possible to reconstruct the signal matrix from measured data, perturbation theory [9] establishes a simple relationship between the singular value of the observed expression matrix $\tilde{X}$ and those of the signal matrix $P$. $P$ is assumed to have finite rank $r$. Its singular value decomposition is thus:

$$P = \sum_{i=1}^{r} \theta_i u_i v_i^T, \quad \text{where } r \ll N, M$$

For scRNA-seq data, we only have to consider singular values $\theta_i > 0$, which means that $\tilde{X}$ potentially has singular values above the MP distribution. Thus, we only need to consider the largest singular values of $\tilde{X}$.

**Theorem 3 (Largest Singular Value for MP)** *([9]) The $r$ largest singular values $\gamma_i(\tilde{X})$ of the $M \times N$ perturbed matrix $\tilde{X}$ exhibit the following behaviour as $M, N \to \infty$ and $\frac{N}{M} \to c$: For each fixed $1 \leq i \leq r$,*

$$\gamma_i(\tilde{X}) \xrightarrow{a.s.} \begin{cases} \sqrt{\frac{(1+\theta_i^2)(c+\theta_i^2)}{\theta_i^2}} & \text{if } i \leq r \text{ and } \theta_i > c^{1/4}, \\ b & \text{otherwise} \end{cases} \tag{9}$$

*Moreover, for each fixed $i > r$, we have that $\gamma_i\left(\tilde{X}_n\right) \xrightarrow{a.s.} b$.*

This theorem establishes a functional relationship between the largest singular values $\gamma_i$ of the measured expression matrix and the singular values $\theta_i$ of the signal matrix $P$. Note that if $\theta_i$ is smaller than or equal to $c^{\frac{1}{4}}$, the corresponding $\gamma_i$ will be equal to $b$, which is the upper limit of the MP distribution. In other words, if the perturbation (signal) is too small, the singular value spectrum of the observed expression matrix $\tilde{X}$ will be just the MP distribution and hence, no meaningful signal can be extracted.

From the above formula we are able to calculate the singular values of the perturbation matrix $P$. These are the values that describe the actual variances of the signal matrix without any contribution of the noise. This is achieved by calculating the inverse function

$$\theta_i(\gamma_i) \xrightarrow{\text{a.s.}} \begin{cases} \sqrt{\frac{2c}{\gamma_i^2-(c+1)-\sqrt{(\gamma_i^2-(c+1))^2-4c}}} & \text{if } \gamma_i > b, \\ c^{\frac{1}{4}} & \text{otherwise} \end{cases} \tag{10}$$

##### 2.4.1 SIGMA

Next, we want to establish how the singular vectors of $\tilde{X}$ depend on the perturbation $P$. In section 2.3 it is described that the elements of the singular vectors will follow a Gaussian distribution for a random matrix and large $N$. The elements of the singular vectors of the perturbation $P$ are deterministic and correspond to biological variance. The following theorem describes the scalar product between the singular vector of the perturbation $P$ and the perturbed matrix $\tilde{X}$.

**Theorem 4 (Norm of Projection of Largest Singular Vectors for MP)** *([9]) Let $\tilde{v}$ the right unit singular vectors of $\tilde{X}$. Then, the norm of projection of the right singular vector is given by*

$$|\langle \tilde{v}_i, v_i \rangle|^2 \xrightarrow{a.s.} \begin{cases} 1 - \frac{c(1+\theta_i^2)}{\theta_i^2(\theta_i^2+c)} & \text{if } \theta_i \geq c^{1/4} \\ 0 & \text{otherwise} \end{cases} \tag{11}$$

This theorem shows the same qualitative behavior as equation 9. If the singular value $\theta_i$ of the perturbation matrix is below the threshold of $c^{\frac{1}{4}}$, the scalar product is zero, indicating that the perturbed matrix $\tilde{X}$ has no relationship to the perturbation $P$. In other words, no relevant signal can be extracted. In the other limit, when the scalar product goes to 1, the singular vectors of the perturbation $P$ are perfectly aligned with the singular vectors of the perturbed matrix $\tilde{X}$. Thus, random noise has a negligible influence on the signal.

The scalar product given by $|\langle \tilde{v}_i, v_i \rangle|^2$ is identical to the squared cosine of the angle between the vectors:

$$\sigma = cos(\alpha)^2 = \left(\frac{\tilde{v} \cdot v}{\|v\| \ \|\tilde{v}\|}\right)^2 = (\tilde{v} \cdot v)^2 = |\langle \tilde{v}_i, v_i \rangle|^2.$$

This holds because the singular vectors are assumed to have norm 1.

We propose $\sigma$ (SIGMA) as a measure of clusterability in scRNA-seq data. If, for a given cluster, there are no values above the MP distribution the signal of the perturbation matrix $P$ can not be recognized any more and $\sigma$ will be zero. If there are singular values above the MP distribution, $\sigma$ evaluates how closely related the singular vectors of the expression matrix $\tilde{X}$ are to those of the perturbation matrix $P$.
We obtain a value of $\sigma$ for each singular value that can be found above the MP distribution. Each of them indicates the signal-to-noise ratio for the variance that the corresponding singular vector explains. Thus, the more singular values are above the MP distribution, the more variances can be found in the data and it can be interpreted as proportional to the number of clusters. In the definition of SIGMA, we have decided to use the maximum of all angles, thus indicating the maximal clusterability that can be achieved from clustering.

##### 2.4.2 G-SIGMA

In accordance with the above definition of SIGMA (2.4.1), we can also define the clusterability, or signal-to-noise ratio, for the gene space. The following theorem describes the equation.

**Theorem 5 (Norm of Projection of Largest Singular Vectors for MP)** *([9]) Let $\tilde{u}$ be the left unit singular vectors of $\tilde{X}$. Then, the norm of projection of the left singular vector by*

$$|\langle \tilde{u_i}, u_i \rangle|^2 \xrightarrow{a.s.} \begin{cases} 1 - \frac{(c+\theta_i^2)}{\theta_i^2(\theta_i^2+1)} & \text{if } \theta_i \geq c^{1/4} \\ 0 & \text{otherwise} \end{cases} \tag{12}$$

For the gene singular vector, $\tilde{\sigma}$ (G-SIGMA) indicates how closely the variance among genes is related to the original variance in the perturbation matrix $P$. For each singular vector, the variance-driving genes correspond to those with the highest absolute loading in the corresponding gene singular vector. Cells with high positive or negative entries in the singular vector have high expression of genes with large positive or negative entries in the corresponding gene singular vector, respectively. This relationship is not a replacement for the calculation of differentially expressed genes, but merely indicates the genes that drive the variance across cells for each singular vector. Based on the value of G-SIGMA, it is possible to evaluate how accurate the determination of differentially expressed genes will be. With a low signal-to-noise ratio, it is more likely to obtain genes differentially expressed that can be attributed to noise. As well as for SIGMA, we obtain several angles, one for each singular value above the MP distribution. Thus, genes driving the variances in gene singular vectors with a higher G-SIGMA are more accurate. We decided, to be consistent, to define G-SIGMA as the highest squared cosine of the angle.

#### 2.5 Clusterability

##### 2.5.1 Assessing clustering quality

We use two different methods to assess clustering quality, the adjusted rand index (ARI) and the silhouette coefficient.
Assuming two partitions, $\mathcal{A}$ and $\mathcal{B}$, of a set of $N$ cells, the rand index is defined as [10]:

$$RI(\mathcal{A}, \mathcal{B}) = \frac{N_{11} + N_{00}}{\binom{N}{2}},$$

where $N_{11}$ is the number of pairs of elements that are in the same cluster in $\mathcal{A}$ and in the same cluster in $\mathcal{B}$. $N_{00}$ is the number of pairs of elements that are in a different cluster in $\mathcal{A}$ and in a different cluster in $\mathcal{B}$. The rand index takes values between 0 and 1, where 0 indicates the complete lack of agreement between the partitions and 1 would indicate identical partitions. Even a random clustering of elements produces a non-zero rand index. The ARI is defined in such a way, that its value is on average 0 for a pair of partitions with randomly permuted cluster labels. A positive ARI thus indicates that partitions agree more than expected to happen by random chance. Let partition $\mathcal{A}$ have $K_{\mathcal{A}}$ clusters of sizes $a_i$ and partition $\mathcal{B}$ have $K_{\mathcal{B}}$ clusters of sizes $b_j$, then the adjusted rand index is defined as:

$$ARI(\mathcal{A},\mathcal{B}) = \frac{RI(\mathcal{A},\mathcal{B}) - \mathbb{E}[RI(\mathcal{A},\mathcal{B})]}{1.0 - \mathbb{E}[RI(\mathcal{A},\mathcal{B})]} = \frac{\binom{N}{2}\sum_{k,m=1}^{K_{\mathcal{A}}K_{\mathcal{B}}}\binom{n_{km}}{2} - \sum_{m=1}^{K_{\mathcal{A}}}\binom{a_k}{2}\sum_{m=1}^{K_{\mathcal{B}}}\binom{b_m}{2}}{\frac{1}{2}\binom{N}{2}\left[\sum_{k=1}^{K_{\mathcal{A}}}\binom{a_k}{2}\sum_{m=1}^{K_{\mathcal{B}}}\binom{b_m}{2}\right] - \sum_{k=1}^{K_{\mathcal{A}}}\binom{a_k}{2}\sum_{m=1}^{K_{\mathcal{B}}}\binom{b_m}{2}}$$

For synthetic data, we take a high ARI between a clustering and the ground truth partition to indicate a clustering of high quality.

Another useful measure for clustering quality is the silhouette coefficient. Let $a(i)$ be the mean distance from point $i$ to all other data points in the same cluster and $b(i)$ be the mean distance from point $i$ to all other points from different clusters, then the silhouette coefficient is defined as [11]:

$$s(i) = \frac{b(i) - a(i)}{max\{a(i), b(i)\}}.$$

For the calculation of the distance, we consider the euclidean distance metric in the space spanned by the singular vectors that are associated with singular values above the MP distribution of the expression matrix $\tilde{X}$ (see 2.3). The final silhouette coefficient is taken as the mean value over all data points. For the calculation of the silhouette coefficient we use the *cluster R package (V 2.1.0)*.

##### 2.5.2 Theoretically achievable clustering quality

A perfect clustering would coincide with the ground truth and obtain an ARI of 1. Here we argue that such a perfect clustering is in general not achievable, if there is noise in the data. In other words there is always a finite Bayes error rate (also called irreducible error) for assigning cells to the appropriate cluster. To construct a Bayes classifier, which achieves the minimal error rate, we need to know the ground truth partition. Hence, we use simulated data. For each ground truth cluster, we fit a multidimensional Gaussian to the elements of the singular vectors of the expression matrix $\tilde{X}$ that correspond to the cells in the respective cluster (see Figure S2a). We only consider singular vectors with singular values above the MP distribution. For the fit we use the *mclust R package (V 5.4.6)*. We then construct a classifier by assigning a cell to the cluster for which it has the highest value of the fitted Gaussian distribution. This corresponds to the best clustering one can achieve if the ground truth partition is known. We define the theoretically achievable adjusted rand index (tARI) as the ARI between this best achievable clustering and the ground truth partition. Similarly, we define the theoretically achievable silhouette coefficient (tSIL) as the silhouette coefficient of the best achievable clustering. Since we use the fitted Gaussian distributions instead of the actual (unknown) distribution of singular vector elements, the constructed classifier only approximates the Bayes classifier. However, we confirmed empirically, that the tARI defined above is an upper bound for all tested clustering methods, which comprises the currently most popular tools used for single-cell RNA-seq data [Figure S2 b, c].
The tARI embodies our notion of clusterability. We define high clusterability as a low Bayes error rate for cluster assignments, which corresponds to a high tARI. We show empirically that our clusterability measure is a proxy of the tARI and thus a way to assess clusterability without knowing the ground truth [Figure 1d].

### 3 Application to single-cell RNA-seq data

#### 3.1 Preprocessing of scRNA-seq data

In the following the necessary preprocessing steps for the application of the clusterability measure for scRNA-seq data are described.

**Transcriptome Mode**
The largest eigenvalue $\lambda_1$ of an expression matrix is typically much larger than all the other singular values and its corresponding singular vector has entries of equal sign, which often have similar magnitude (of order $\frac{1}{\sqrt{N}}$, which is the ideal value in the perfectly homogeneous case). This singular vector reflects a general, global trend in the data. This structure has been observed for many empirical data matrices. (In time series analysis of the stock market, this singular vector is called the "market mode" since it corresponds to a trend that is common across many stocks [3]). Here, we refer to this singular vector as "transcriptome mode" since it reflects a trend that is shared across the whole transcriptome (see Figure S1). In order to reduce the influence of this singular value on the calculation of the MP fit, we center the expression matrix $\tilde{X}$ gene-wise. As a result, the singular value of the transcriptome mode will be reduced to a value close to 0.

**Normalization**
The efficiency of the capture of transcripts and their conversion to cDNA is known to be highly variable between cells. Hence, single-cell gene expression data is usually normalized cell-wise. We have tested several normalization methods but none of them seemed sufficient to remove all technical variability in the data. Thus, in section 3.2 we describe a method to reduce these effects for our clusterability measure $\sigma$. Nevertheless, we normalize the expression to the total counts per cell and subsequently log-transform to stabilize the variance.

**Gene distribution**
Gene expression is typically modelled by a Poisson, negative binomial or zero inflated negative binomial distribution. However, the parameters of these distributions differ between genes, this violates the assumptions of the MP theorem, where all values are sampled from the same distribution. In practice, gene-wise standardization to a mean of 0 and standard deviation of 1 mostly circumvents this problem. Additionally, we have observed that there is a bias resulting from variations in cells. These biases are as well reduced by standardising the cells to a mean of 0 and standard deviation of 1 (see Figure S1 c,d). This is equivalent to calculating the eigenvalues and vectors of a correlation matrix instead of a covariance matrix.

**Zero inflation**
Another factor to be considered is the large amount of zero values in scRNA-seq data. These zeros might be on the one hand due to technical artefacts (low efficiency, dropout) or simply due to low, stochastic gene expression. After performing the above mentioned preprocessing steps we mostly do not observe deviations from the MP distribution. However, this is a known problem discussed within the framework of sparsity induced singular values. For single cell RNA-seq data an extensive analysis has been performed in [8], where the authors observe deviations from the MP distribution caused by sparsity. The authors suggest the exclusion of outlier genes that can be identified through the fit of the MP distribution. For SIGMA we do not use this preprocessing step, however we do exclude genes that have a high expression in only a few number of cells.

##### 3.2 Regressing out unwanted sources of variability (Confounder Regression)

scRNA-seq data suffers from several sources of technical variability that can obscure or even be mistaken for relevant biological signal. One of the most important of these is the variable efficiency of mRNA capture and cDNA conversion. The total number of detected transcripts per cell is typically taken as a proxy of this efficiency. There are also biological processes that can cause unwanted signal. Most cells are stressed due to the tissue dissociation necessary for single-cell library preparation. The percentage of expression coming from mitochondrial genes or the expression of marker genes for stress can be used to estimate the level of stress. Different metabolic states of cells might be reflected in the level of ribosomal gene expression and many genes fluctuate with the cell cycle. Here, we seek to establish a method to remove any effect of these nuisance variables on the clusterability measure.

We model the signal matrix $P$ as a sum of relevant signal $B$ and unwanted signal due to nuisance variables $Y$. Inspired by published approaches to expression data normalization [12, 13], we model the influence of $Y$ by linear regression. This is a valid approach because the regression is performed on the singular vectors of $\tilde{X}$, which contain Gaussian distributed noise. Given the singular value decomposition of $\tilde{X}$ and singular vectors $\tilde{v}_i$,

$$\tilde{v}_i = \beta Z, \qquad \text{with } \beta \in \mathbb{R}^k \tag{13}$$

where $Z \in \mathbb{R}^{N \times k}$ is a matrix of covariates, such as the total counts per cell, with $k$ the number of covariates and $N$ the number of cells. Each covariate is normalized to 1 such that the range agrees with the range of the singular vectors. The amount of variance explained by the nuisance parameters is then given by the value of the adjusted R squared ($R^2_{adj}$) of this linear regression. Since the eigenvalues of the cell-cell correlation matrix can be interpreted as the amount of variance explained, we reduce the eigenvalues $\lambda_i$ by $\tilde{\lambda}_i = (1 - |R^2_{adj}|)\lambda_i$. In the next step, we calculate adjusted singular values by $\tilde{\gamma}_i = \sqrt{\tilde{\lambda}_i}$ and use these adjusted singular values $\tilde{\gamma}_i$ for the consecutive steps in the calculation of the clusterability measure.

##### 3.3 Algorithm

The procedure to obtain the clusterability measure involves the following steps:

1. Preprocess the single cell expression matrix as described in section 3.1:
   (a) Normalization
   (b) Log-transformation
   (c) Standardization gene-wise
   (d) Standardization cell-wise
2. Calculate the singular value decomposition of the gene expression matrix $\tilde{X}$.
3. Fit the MP distribution to the singular values (equation 5).
4. Determine singular values/vectors that correspond to non-random variability using the Tracy-Widom distribution (equation 7) or the Shapiro-Wilk test (equation 8), respectively.
5. Adjust the singular values for effects of nuisance variables by linear regression (equation 13).
6. Calculate the singular values $\theta_i$ of the signal matrix $P$ using the inverse of equation 9, given by 10.
7. Calculate the projections of the singular vectors of the expression matrix $\tilde{X}$ on the corresponding singular vector of the signal matrix $P$ with equations 11 for the singular vectors and 12 for the gene singular vectors.
8. The clusterability measure is the largest of the projections for the singular vectors obtained in the previous step.

#### 4 References

[1] O Bohigas, M J Giannoni, and C Schmit. Characterization of chaotic quantum spectra and universality of level fluctuation laws. *Physical Review Letters*, 52, 1984.

[2] Marc Potters, Jean-Philippe Bouchaud, and Laurent Laloux. Financial applications of random matrix theory: Old laces and new pieces. *Science & Finance, Capital Fund Management, Science & Finance (CFM) working paper archive*, 36, 2005.

[3] Mel Macmahon and Diego Garlaschelli. Community detection for correlation matrices. *Physcial Review*, 021006:1–34, 2015.

[4] Joël Bun, Jean philippe Bouchaud, and Marc Potters. Cleaning large correlation matrices : Tools from random matrix theory. *Physics Reports*, 666:1–109, 2017.

[5] Eugene P Wigner. Characteristic vectors of bordered matrices with infinite dimensions. *Source: Annals of Mathematics*, 62:548–564, 1955.

[6] Craig A. Tracy and Harold Widom. Level-spacing distributions and the airy kernel. *Communications in Mathematical Physics*, 159(1):151–174, 1994.

[7] Giacomo Livan, Marcel Novaes, and Pierpaolo Vivo. *Introduction to Random Matrices - Theory and Practice*, volume 26. Springer, 12 2017.

[8] Luis Aparicio, Mykola Bordyuh, Andrew J Blumberg, and Raul Rabadan. Article a random matrix theory approach to denoise single-cell data a random matrix theory approach to denoise single-cell data. *Patterns*, page 100035, 2020.

[9] Florent Benaych-georges and Raj Rao. The singular values and vectors of low rank perturbations of large rectangular random matrices. *Journal of Multivariate Analysis*, 111:120–135, 2012.

[10] Alexander J Gates and Yong-Yeol Ahn. The impact of random models on clustering similarity. *Journal of Machine Learning Research*, 18:1–28, 2017.

[11] Peter J Rousseeuw. Silhouettes: a graphical aid to the interpretation and validation of cluster analysis. *Journal of Computational and Applied Mathematics*, 20:53–65, 1987.

[12] Dominic Grün. Revealing dynamics of gene expression variability in cell state space. *Nature Methods*, 17:45–49, 1 2020.

[13] Christoph Hafemeister and Rahul Satija. Normalization and variance stabilization of single-cell rna-seq data using regularized negative binomial regression. *Genome Biology*, 20, 12 2019.

### Supplementary Note 2

Application of SIGMA to our previously published single-cell RNA-sequencing study of the human fetal kidney[1] revealed two distinct groups of clusters (Fig. 3a). Connecting tubule (CnT), nephron progenitor cells-a (NPCa), nephron progenitor cells-b (NPCb), and mesangial cells (Mes) all obtained a SIGMA of 0, which signified that these clusters consist of pure populations with homogeneous gene-expression profiles. The rest of the clusters obtained higher values of SIGMA, indicating that they contained subpopulations that were previously overlooked. We explored all clusters with highest SIGMA and chose to further investigate clusters in which new cell populations were identified: the ureteric bud/collecting duct (UBCD), the S-Shaped Body proximal precursor cells (SSBpr), and the Interstitial cells a (ICa), with SIGMAs of 0.97, 0.95, and 0.93, respectively.

#### UBCD

The analysis of this cluster yielded two clearly separate subpopulations (Fig. 3b, Fig. S8b). The bigger subpopulation contained developing collecting duct cells and their precursors (ureteric bud), indicated by the expression of genes such as WFDC2, AQP2, CLDN3, MMP7, and CALB1. In contrast, the smaller sub-cluster showed little or no expression of the aforementioned genes and was characterized by UPK1A and UPK1B, well-known markers of the urothelial epithelium, which constitutes the inner lining of the ureter. The presence of such cells in our data is plausible given that the whole fetal kidney was used in our sequencing experiment. Both DE analysis (Supplementary Table 2) and inspection of the top variance-driving genes (Fig. S8e) revealed SPINK1, UPK2, S100A6, KRT7, and KRT19 as additional markers. Staining of week 15 fetal kidney sections with UPK1A and KRT7 antibodies confirmed our interpretation (Fig. 3c, Fig. S9a). UPK1A was restricted to the superficial urothelial cells in major and minor calyces as well as the developing ureter. KRT7 was expressed more broadly, across the superficial, intermediate, and basal urothelium. Both KRT7 and UPK1A were completely absent from the whole collecting system and the branching ureteric bud, marked by CDH1 (Fig. S9a).

#### SSBpr

Sub-clustering the SSBpr population showed the presence of 3 subpopulations (Fig. 3b, Fig. S8b). One subpopulation contained markers of proximal cell precursors (GPC3, LHX1, CADM2) together with low expression of AMN and APOE (see Supplementary Table 2), which is consistent with the original annotation of the cluster.

A second subpopulation, contiguous to the previous one, showed the expression of CLDN1, which is expressed in the proximal epithelium, together with CITED2, expressed in developing podocytes. This suggested parietal epithelial cells (PECs) as the most likely cell type, as these cells were reported to share several markers with both proximal epithelium and podocytes[2]. To confirm this interpretation, we performed Immunostaining of CLDN1, as well as CAV2 and AKAP12 which were found by DE analysis (Fig. 3d, Fig. S9b). Interestingly, CLDN1 was found in all segments of the S-shaped body except in the precursors of the PECs, which are the thin layer of cells at the lateral side of the proximal segment of the SSB. CLDN1 appeared in the parietal epithelium only at the capillary loop stage and continued to be expressed in all PECs in more mature glomeruli. CAV2 was present in the parietal epithelium in developing glomeruli, but also in the endothelial cells of both the glomerular capillaries and the surrounding vasculature. Intriguingly, CAV2 overlapped with CLDN1 only in a subpopulation of PECs in individual glomeruli, which might indicate previously unobserved heterogeneity within these cells in the developing kidney. Only AKAP12 marked the precursors of the PECs in S-shaped bodies and continued to be abundantly expressed. However, AKAP12, was not specific to PECs, as it was also expressed in interstitial cells in the cortex.

Finally, a third, small and distinct subpopulation in the SSBpr cluster expressed distal tubule markers (SPP1, ODC1, IRX3, and S100A10), suggesting that these cells were misclassified during the original clustering. This shows that SIGMA can pinpoint clustering errors, making it a useful tool for clustering quality control.

#### ICa

This cluster consisted of 5 subpopulations (Fig. 3b, Fig. S8b). All subpopulations expressed markers of the renal interstitium. One also expressed genes found uniquely expressed in other cell types (EPCAM, CD24, BST2, NNAT, DAPL1) and thus likely contains doublets. Another small subset was characterized by markers of mesangial cells (MGP, ACTA2, PDGFRB), suggesting that it contains mesangial cells erroneously grouped with the ICa or renal pericytes, which share a similar gene expression profile[3]. Another subpopulation showed high expression of non-specific genes related to components and regulators of microtubules together with metabolic, mitochondrial and stress-related genes (H2AFZ, TUBA1B, TYMS, STMN1, DUT, MT-CO3, MT-ND5).

The two remaining subpopulations were clearly interstitial but their gene expression profiles could not be linked to known interstitial populations, likely due to the dearth of knowledge about the renal stroma. We hypothesized that these two subpopulations were localized in different regions of the kidney. To test this idea, we stained fetal kidney sections with POSTN, CLDN11, and SULT1E1 (Fig. 3e, Fig. S9c), which were identified by DE analysis and inspection of the top variance-driving genes. SULT1E1 was highly expressed in the pelvic area in the immediate vicinity of the developing ureter, as well as the inner and outer medulla, preferentially surrounding tubules. This marker might thus indicate the medullary interstitium as well as pelvic smooth muscle cells. Staining with CLDN11 showed a higher signal in the medulla and papilla, similar to SULT1E1, but with a wider spatial distribution. In contrast to SULT1E1, CLDN11 was also expressed in groups of cortical interstitial cells, situated directly underneath the renal capsule, in the nephrogenic zone. CLDN11 might thus also be expressed by the interstitial progenitor cells or their immediate progeny. Lastly, POSTN was mainly found in the renal cortex surrounding tubules and glomerular microvasculature. POSTN was also expressed in cortical blood vessels with larger diameters together with their arborizations. POSTN is a secreted extracellular matrix protein known to be expressed in cardiac smooth muscle cells, as well as connective tissues. Here, POSTN might mark smooth muscle cells of the cortical vasculature.

In conclusion, a reanalysis of our previously published data showed the ability of SIGMA to reveal overlooked subpopulations. Interestingly, SIGMA identified sub-clusters with only a few cells (41 developing PECs, 68 urothelial cells, 29 distal cells), highlighting its sensitivity to relevant substructure hidden within a bigger cluster. Finally, SIGMA was also useful to pinpoint clustering errors and the presence of doublets, which makes it useful for quality control prior to DE analysis.

| Data Set | Data description | Reference | Data download | Simulation parameters | Normalization Method | Protocol |
|---|---|---|---|---|---|---|
| Real scRNA-seq data sets | | | | | | |
| BMNC | Bone marrow mononuclear cells, one human donor | https://www.cell.com/cell/fulltext/S0092-8674(19)30559-8 | R package *SeuratData (V 0.2.1)*, "bmcite" | - | *Seurat (V 3.1.5):* NormalizeData | 10X Chromium |
| fetal kidney | fetal kidney from W16 of gestation | https://journals.plos.org/plosbiology/article?id=10.1371/journal.pbio.3000152 | - | - | R package *scran* by original authors | 10X Chromium |
| PBMC | Peripheral Blood Mononuclear Cells | https://support.10xgenomics.com/single-cell-gene-expression/datasets/1.1.0/pbmc3k? | https://cf.10xgenomics.com/samples/cell/pbmc3k/pbmc3k_filtered_gene_bc_matrices.tar.gz | - | *scanpy (V 1.4.6)* and seurat (V 3.9.9.9010)<br>As described in the respective pipeline. | 10X Chromium |
| RNA-mix | RNA was extracted from 3 cell lines (HCC827, H1975 and H2228) and mixed in different proportions. | https://www.nature.com/articles/s41592-019-0425-8 | https://github.com/LuyiTian/sc_mixology | - | *scran (V 1.14.6)*: quickCluster, computeSumFactors | CEL-seq2 |
| Simulated data sets | | | | | | |
| Fig. 1b | Simulated data sets with 5 clusters | - | - | High signal-to-noise:<br>nGenes = 1000, batchCells = 500, group.prob =c(0.03809524, 0.11904762, 0.20000000, 0.28095238, 0.36190476), path.nonlinearProb = 0, out.prob = 0.1, dropout.type = "none", lib.loc = 11, de.downProb = 0.5, de.prob = 0.1, seed = 3, de.facLoc = 0.1<br>Low signal-to-noise:<br>nGenes = 1000, batchCells = 500, group.prob = c(0.03809524, 0.11904762, 0.20000000, 0.28095238, 0.36190476), path.nonlinearProb = 0, out.prob = 0.1, dropout.type = "none", lib.loc = 11, de.downProb = 0.5, de.prob = 0.03, seed = 3, de.facLoc = 0.1 | Dividing by the total number of transcripts and multiplying by 10000 | - |
| Fig. 1c, Extended Data Fig. 6a | Simulated data sets with 5 clusters | - | - | Parameters:<br>nGenes = 1000, batchCells = N, group.prob = c(0.03809524, 0.11904762, 0.20000000, 0.28095238, 0.36190476), path.nonlinearProb = 0, out.prob = 0.1, dropout.type = "none", lib.loc = 11, de.downProb = 0.5, de.prob = seq(0, 0.15, length.out = 50), seed = 3, de.facLoc = 0.1<br>3 simulations:<br>For N equal to 500, 1000 and 2000 | Dividing by the total number of transcripts and multiplying by 10000 | - |
| Fig. 1d, Extended Data Fig. 2bc, Extended Data Fig. 3a-c | Simulated data sets with 2 clusters | - | - | Varying DE genes:<br>nGenes = 1000, batchCells = 500, group.prob = c(0.3, 0.7), path.nonlinearProb = 0, out.prob = 0.1, dropout.type = "none", lib.loc = 11, de.downProb = 0.5, de.prob = seq(0, 0.15, length.out = 50), seed = seed, de.facLoc = 0.1<br>Varying FC:<br>nGenes = 1000, batchCells = 500, group.prob = c(0.3, 0.7), path.nonlinearProb = 0, out.prob = 0.1, dropout.type = "none", lib.loc = 11, de.downProb = 0.5, de.prob = 0.05, seed = seed, de.facLoc = seq(0, 0.3, length.out = 50), de.facScale = 0.1 | Dividing by the total number of transcripts and multiplying by 10000 | - |
| Extended Data Fig. 6 b-d | Simulated data set with 3 clusters | - | - | Parameters:<br>nGenes = 1000, batchCells = 500, group.prob = c(0.65, 0.2, 0.15), path.nonlinearProb = 0, out.prob = 0.1, dropout.type = "none", lib.loc = 11, de.downProb = 0.5, de.prob = 0.1, seed = 69, de.facLoc = 0.1 | Dividing by the total number of transcripts and multiplying by 10000 | - |
